# ediTONSIL: Activation-Neutral Gene Editing of CD4 T cells in Human ex Vivo Tonsil Cultures

**DOI:** 10.1101/2023.08.21.554089

**Authors:** K. Morath, L. Sadhu, G. Dyckhoff, Madeleine Gapp, Oliver T. Keppler, O.T. Fackler

## Abstract

**Motivation:** CD4 T cells are central players of adaptive immunity that rapidly change activation states in response to exogenous T cell receptor (TCR) stimulation. While molecular processes in activated human CD4 T cells from peripheral blood are well studied, resting CD4 T cells are refractory to gene-editing transduction and transfection methods without prior activation. Knowledge on the molecular biology of truly resting CD4 T cells is therefore lacking. We present here the culture and editing workflow ediTONSIL that allows for gene editing of tissue-resident CD4 T cells without compromising their activation state or immunological function. This will enable molecular mechanistic analyses of this cell type in their native/physiological tissue context.

**Summary:** The molecular and immunological properties of resting CD4 T cell biology are understudied due to the lack of suitable gene editing methods. Here we describe the *ex vivo* culture and gene editing methodology ediTONSIL for tissue-resident CD4 T cells from human tonsil tissue. CRISPR/Cas9 RNP nucleofection under optimized culture conditions and cytokine concentrations results in knock out efficacies of over 90%, which are e.g. sufficient to prevent HIV-1 infection when targeting the viral entry co-receptor CXCR4. EdiTONSIL does not impair tonsil CD4 T cell viability, activation state or immunocompetence and does not require exogenous activation. Editing can be performed on multiple cell types in bulk cultures or of CD4 T cells previously isolated from tonsils that can be added back to the non-CD4 T cell fraction post gene editing to reassemble into immunocompetent organotypic lymphoid aggregate structures. This highly efficient and versatile workflow for gene editing of tonsillar CD4 T cells enables the dissection of molecular mechanisms in *ex vivo* cultures of human lymphoid tissue and can be adapted to other tonsil-resident cell types.

**Graphical Abstract:** 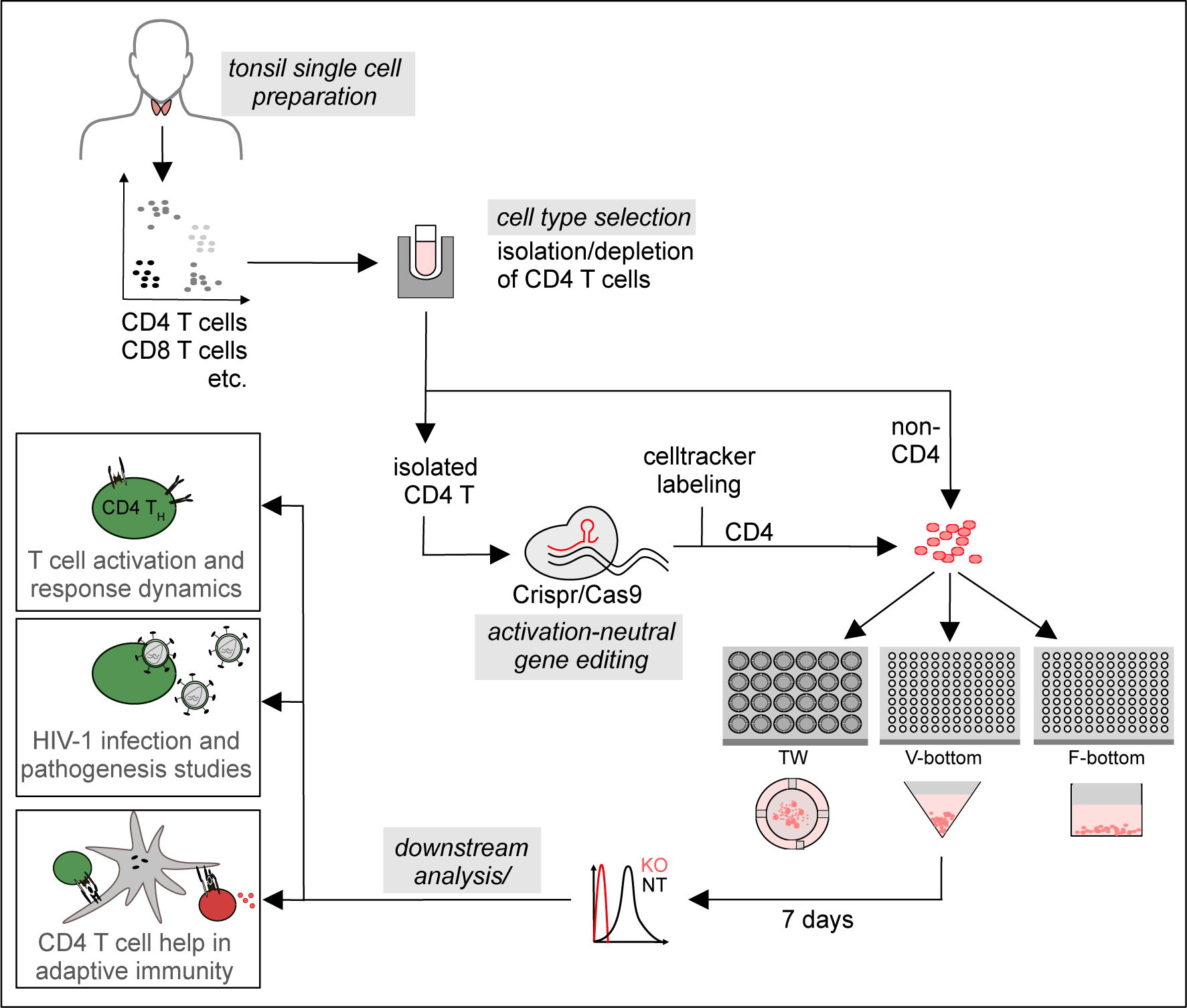

## Introduction

Adaptive immune reactions involve fine-tuned, complex and dynamic interactions between different cell types that define the remarkable potency of our immune system. In the case of CD4 T cells, interactions with antigen-presenting cells (APCs) that display MHC-II-restricted specific antigenic peptides in the context of close cell-cell conjugates (immunological synapse) drive CD4 T cell activation, proliferation and differentiation to elicit effector function that include helper functions to B and CD8 T cells towards humoral and cytotoxic immune responses, respectively^1^. These APC-effector interactions take place in lymphoid organs and tissue architecture, composition and density are essential parameters for efficacy and outcome of these reactions. Despite this essential impact of the tissue environment, most studies on CD4 T cell function employ CD4 T cells from readily available peripheral blood, which display activation and differentiation states that are markedly distinct from that of tissue-resident CD4 T cells. An alternative approach is the use of small animal models (mostly mouse), in which complex immune reactions can be studied. While these models provide fundamental insight, they also have important limitations with respect to human pathogenesis as they differ in tissue composition, the spectrum of cell surface markers to identify cell populations, life span and exposure to a complex immune-stimulatory environment^2^. These differences are particularly relevant for studies of pathogen-host interactions such as HIV infection as even transgenic or humanized mouse models do not fully mirror the complex pathogenesis induced by the infection in humans^3,4^. To overcome this limitation, organotypic *ex vivo* cultures of lymphoid tissue explants were developed and cultures of human tissue organ proved most useful. Grown as tissue blocks (human lymphoid histoculture, HLH)^5–12^ and in suspension (human lymphoid aggregate culture, HLAC)^9,10,13–16^, human tonsil tissue is permissive to HIV-1 infection and mirrors hallmarks of HIV pathogenesis such as CD4 T cell depletion. Of note, the original version of HLH retained the ability to elicit adaptive antibody responses to recall antigen^7,17,18^ and a recent optimization of cell purification and reassembly also improved the immunological features of HLACs, allowing the testing of vaccine candidates^19,20^.

The key limitation in this organotypic and immunocompetent cell culture system is the lack of methodology for targeted gene-editing of specific cell types, which currently precludes detailed molecular dissection of cell function in this tissue-like context. Most available methods for gene editing of primary human lymphocytes require prior cell activation for efficient delivery of viral vectors, expression of transgenes or sufficiently high transfection rates^21–23^. These approaches are typically associated with high cytotoxicity, editing rates that require subsequent isolation of edited cells, and are limited to the analysis to the second response of cells to a stimulus that had returned to a phenotypically resting state after a first round of activation. We recently introduced a CRISPR/Cas9 RNP nucleofection approach for peripheral blood CD4 T cells that resulted in highly efficient gene editing without compromising the resting state of these cells^21^. In this study, we adapted this protocol to tissue-resident cells in *ex vivo* tonsil cultures that are first characterized with respect to different culture set-ups and define the ediTONSIL workflow that enables the activation neutral editing of bulk and isolated cell population for the molecular dissection of complex immune cell interaction in these organotypic cultures.

## Results

### Optimizing culture conditions of ex vivo tonsil suspension cultures

The viability of *ex vivo* tonsil cultures is currently limited to 2-3 weeks^5,19,24^. To establish optimized conditions, we compared the impact of different culture conditions on cell viability, activation/differentiation as well as response to stimulation. Tonsil tissue from healthy donors that underwent routine tonsillectomy or tonsillotomy were dispersed through a cell strainer for subsequent isolation of mononuclear cells (Fig. 1A). At this step, cells can either be plated for immediate culture or frozen in aliquots for future use. For comparison, we plated cells in transwells (TW) or V- or F-bottom 96-well plates for subsequent phenotypic characterization by flow cytometry (Fig. 1A). Cell viability was assessed by staining with the dead-cell dye zombie violet (over 95% viable cells at day 0) and cell type composition was determined by the detection of specific cell surface markers (Fig. 1B). Expectedly, cell viability decreased over time, reaching approx. 20% after 15 days of culture. This decline was independent of the culture format (Fig. 1D). With respect to specific cell populations, the relative amount of T cells slightly increased with time at constant CD4 vs. CD8 T cell ratios at the expense of APCs. The only notable impact of the culture format was a marked enrichment of macrophages among all antigen-presenting cells (APC) in TW cultures (Fig. 1F). Activation of CD4 T cells with staphylococcal enterotoxin B (SEB) superantigen triggered potent cell surface exposure of the activation markers CD69, CD25 and CD38 within seven days, while HLA-DR surface levels remained largely unaltered (Fig. 1G-I, supplementary Fig. 1A-E). While the activation pattern was similar in all three culture conditions, exposure of the early activation marker CD69 was significantly more transient and induction of surface CD38 more pronounced in TW cultures.

**FIG 1.**
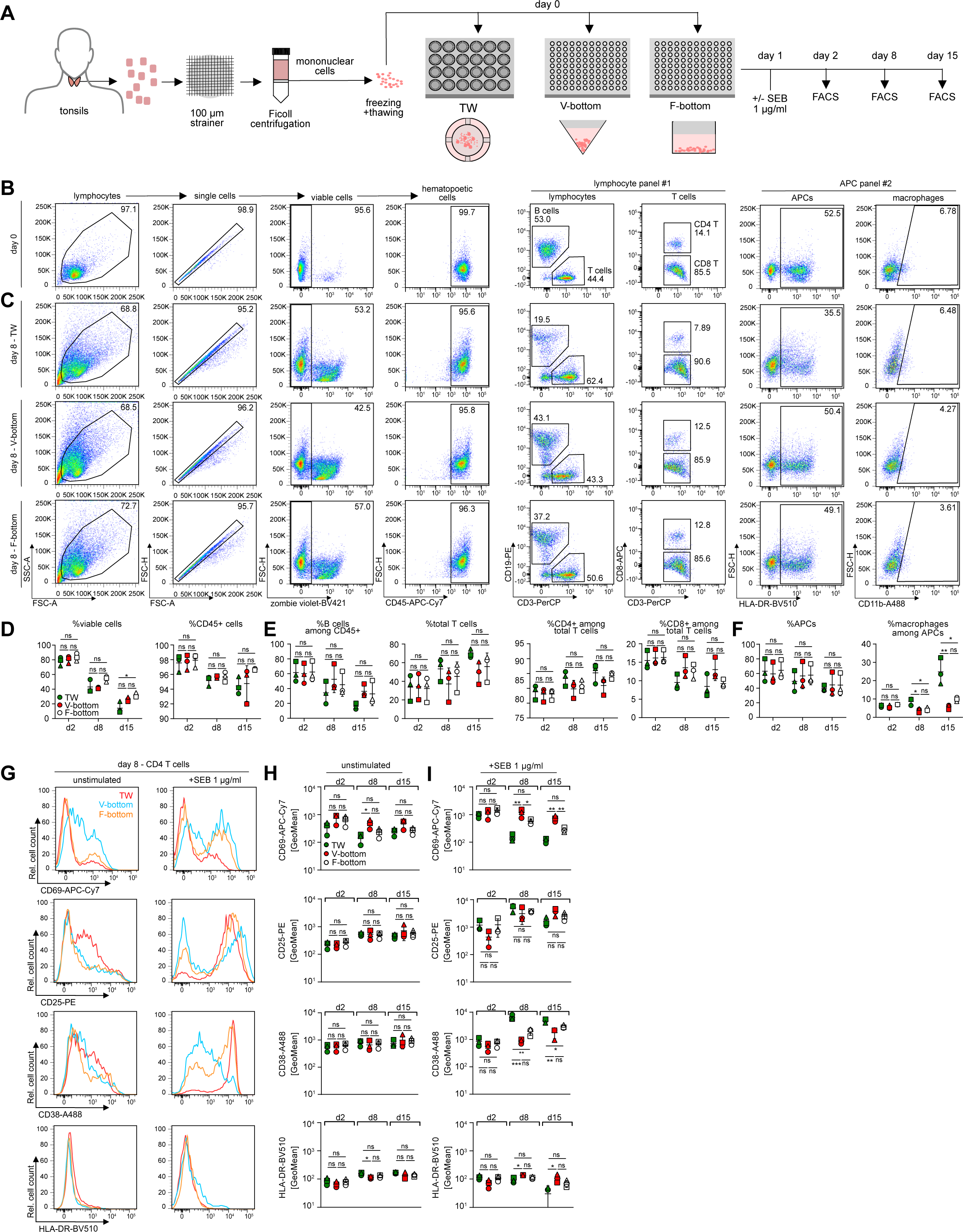
Comparison of cell culture plates for human tonsil cell culture. (A) Schematic of the experimental setup. Cell suspensions were prepared from human tonsils, frozen and thawed for culture in three different cell culture plate systems, 24-well transwell culture (TW), 96-well V-bottom (V-bottom) or 96-well F-bottom (F-bottom) plates at 10x10^6^ cells/ml. Cell composition was analyzed on day 2, day 8 and day 15 post seeding by flow cytometry analysis. Activation response of CD4 T cells in the different setups was measured 1, 7 and 14 days post addition of SEB at 1 µg/ml by flow cytometry. (B) Representative dot plots and gating strategy for identification of cell populations within tonsil bulk cell cultures at day 0, using two antibody cocktails for characterization of lymphoid cells (panel #1) and myeloid cells (panel #2). (C) Representative dot plots at day 8 of culture for TW, V-bottom and F-bottom setups. (D)-(F) Quantification of B and C for three tonsils in TW (green), V-bottom (red) or F-bottom (white) setups. (G) Representative histograms of CD4 T cell surface activation marker expression of CD25, CD69, CD38 and HLA-DR on day 8 of culture with (+SEB 1 µg/ml) or without (unstimulated) exogenous activation. (H) and (I) Quantification of CD69, CD25, CD38 and HLA-DR signal on CD4 T cells without (H) or with (I) exogenous activation as shown in G for three tonsils in TW (green), V-bottom (red) or F-bottom (white) setups. Shown are means with SD. Each symbol represents one donor. Statistical significance was assessed by one-way ANOVA (ns, p>0.05; *, p<0.05; **, p<0.01; ***, p<0.001; ****, p<0.0001).

We next conducted a function characterization of *ex vivo* tonsil cultures under these different conditions. Since HIV-1 typically replicates efficiently in *ex vivo* tonsil cultures and induces the specific depletion of CD4 T cells that is characteristic for AIDS pathogenesis^5,7,25,26^, we first analyzed their permissivity for HIV-1 infection. Following plating, cultures were infected with the full length, replication-competent HIV-1 variant NL4.3SF2Nef one day later and HIV-1 replication and CD4 T cell depletion were quantified over time by assessing the activity of the viral enzyme reverse transcriptase in the cell culture supernatant (SG-PERT assay^27,28^), determining the number of productively infected cells by intracellular staining for the viral capsid protein p24, and defining the ratio of CD4 vs. CD8 T cells by flow cytometry (Fig. 2B-F). As indicated by the reduction in CD4 T cells paralleled by a relative increase in CD8 T cells, CD4 T cell depletion upon HIV-1 infection was observed with comparable magnitude and kinetics in all three culture conditions. The frequency of productively infected, p24^+^ cells reached a peak 3 days post infection (dpi) and then declined, but V-bottom cultures tended to allow the detection of higher frequencies of productively infected cells among viable cells (Fig. 2E). Finally, virus replication in the culture as determined by the SG-PERT assay also rapidly increased in the first dpi and then slowly declined with time but the virus titers observed in TW cultures were markedly lower to those in the other culture conditions (Fig. 2F).

**FIG 2.**
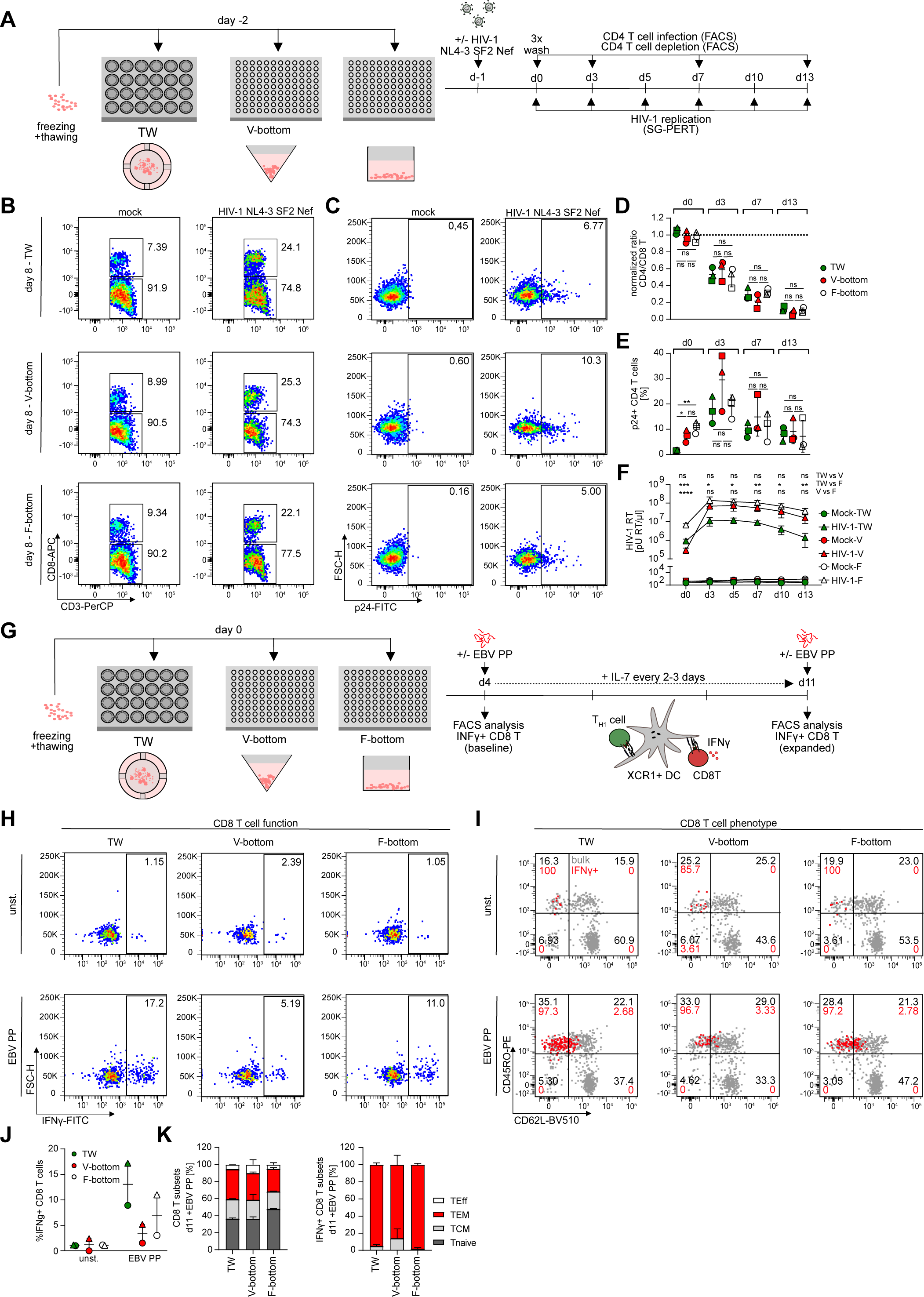
Different culture setups for human tonsil cells support HIV-1 infection to similar extend but show distinct preservation of immunocompetency. (A) Schematic of the experimental setup. Cryopreserved human tonsil cells were thawed and seeded in 24-well transwell (TW), 96-well V-bottom or 96-well F-bottom cell culture plates one day prior to infection with 1.5x10^5^ BCU per 2x10^6^ cells using HIV-1 NL4.3SF2Nef virus. On subsequent days, HIV-1 infection was characterized by analyzing CD4 T cell depletion by a decreased CD4/CD8 T cell ratio and CD4 T cell infection by intracellular flow cytometry staining for HIV-1 capsid protein (p24) and lastly, detection of newly produced HIV-1 virions in the culture supernatant by SG-PERT. (B) Representative dot plots showing CD4 and CD8 T cells subgated from CD3 T cells as well as p24 signal in CD4 T cells for the three culture setups. (D) and (E) Quantification of B and C for three tonsils for the respective harvesting timepoints cultured in TW (green), V-bottom (red) or F-bottom (white) setups. Shown are means with SD. Each symbol represents one donor. (F) SG-PERT analysis of HIV-1 RT activity in the supernatant for mock or HIV-1 infected cultures in TW (green), V-bottom (red) or F-bottom (white) setups. Of note, the higher culture volume in TW plates is a confounding factor for lower RT activity/µl. Shown are means with SD of three tonsils per data point. (G) Schematic of the experimental setup for analysis of EBV-specific memory CD8 T cell reactivation in tonsil cultures. Briefly, cells were thawed and seeded in 24-well transwell (TW) or 96-well V- or F-bottom plates as depicted. On day 4, EBV peptide pool was added together with IL-7 for 7 days. On day 4 and day 11, baseline and expanded EBV-specific CD8 T cells, respectively, were identified by production of IFNγ following stimulation with the EBV peptide pool. (H) Representative dot plots of CD8 T cells in bulk tonsil cell cultures on day 11 of culture with (EBV PP) or without (unstimulated) EBV peptide pool. (I) Characterization of memory subsets based on expression of CD45RO and CD62L among total (grey) or superimposed IFNγ+ (red) CD8 T cells on day 11 following peptide stimulation of tonsil cells cultured with (EBV PP) or without (unstimulated) EBV peptide pool for 7 days. (J) Quantification of (H) for two tonsils per condition. Shown are means with range. (K) Quantification of (I) for total or IFNγ+ CD8 T cells on day 11 in EBV PP-stimulated cultures. T_Eff_ = T effector cells, T_EM_ = effector memory cells, T_CM_ = central memory cells, T_naive_ = naïve T cells. Statistical significance was assessed by one-way ANOVA (ns, p>0.05; *, p<0.05; **, p<0.01; ***, p<0.001; ****, p<0.0001).

To assay the immune competence of these cultures, we probed their ability to mount CD4 T cell help to CD8 T cells in response to presentation of a recall antigen by XCR1^+^ dendritic cells (DC)^29–32^ (Fig. 2G and supplementary Fig. S1F-P). To this end, cultures were treated with a pool of peptides derived from Epstein Barr Virus (EBV), a very common and lifelong infection in the general population that can trigger efficient CD4 and CD8 T cell responses^33–35^. The presence and expansion of reactive cells was then probed one week later by the quantification of interferon gamma (INFγ)-producing CD8 T cells upon peptide restimulation. We observed expansion of INFγ^+^ CD8 T cells in response to EBV peptides in all culture conditions but this effect was clearly most pronounced in TW and F-bottom cultures and V-bottom cultures only allowed for residual expansion of INFγ^+^ CD8 T cells (1.5/5.2% IFNγ+ CD8 T cells for two tonsils compared to above 3.0/8.9 and 11.0/17.2% in F-bottom and TW cultures for the same tonsils, respectively) (Fig. 2H, J). The subset distribution of total or CD8 T INFγ^+^ cells, however, was comparable in all culture conditions (mean of 36.0-47.8% T_naive_, 20.3-23.1% T_CM_, 26.5-35.1% T_EM_ and 5.4-10.5% T_Eff_ cells among total CD8 T cells across the culture set-ups compared to 1.7-13.9% T_CM_ and 86.1-98.4% T_EM_ among IFNγ+ CD8 T cells) (Fig. 2I, K) Of note, prior depletion of CD4 T cells from these cultures abrogated the induction of INFγ^+^ CD8 T cells, indicating that CD4 T cell help is critical for this expansion (supplementary Fig. S1M-P). The culturing format thus has important implications for the functional properties of these cultures with all three culture systems supporting HIV-1 replication yet superior preservation of immunocompetence in TW and F-bottom cultures, which are therefore more suitable for the analysis of complex immune reactions.

### Establishment of ediTONSIL gene knockout in bulk tonsil cultures

We next attempted to perform gene editing of tonsil cells, here termed ediTONSIL, by example of the chemokine receptor CXCR4 that serves as an essential co-receptor for entry of X4-tropic HIV-1 strains and is abundantly expressed on all major immune cell populations. To this end, thawed bulk tonsil cells were nucleofected with CRISPR/Cas9 RNP complex and cultured for 7 days. Knockout (KO) was assessed by flow cytometry which enabled the analysis of KO efficiency in distinct cell (sub)populations and differentiation states (Fig. 3A, B and G). Among the lymphocyte populations, B cells, CD4 and CD8 T cells displayed highly efficient KO of CXCR4 compared to a non-targeting (NT) RNP control within 7 days (Fig. 3C-F; B cells CXCR4 surface level mean for NT 9793.0±1338.7 arbitrary units (AU) vs. KO 166.3±41.7, CD4 T cells 2996.3±341.8 vs. 171.3±26.8 and CD8 T cells 2321.0±328.4 vs. 103.6±31.4) irrespective of the memory differentiation state of CD4 or CD8 T cells (Fig. 3G-M; CD4 T T_CM_ CXCR4 surface level mean for NT 5977.0±1556.3 AU vs. KO 114.3±3.2, T_naive_ 9543.7±1419.5 vs. 230.3±66.0 and T_EM_ 8065.0±1392.8 vs. 173.0±22.3; CD8 T T_CM_ 5051.3±1527.6 vs. 99.9±8.1, T_naive_ 4606.3±2177.5 vs. 137.7±14.4 and T_EM_ 4049.0±765.2 vs. 117.8±70.4) and KO efficiency did not require further exogenous stimulation by SEB (supplementary Fig. S2A-E). Among myeloid cells, macrophages and dendritic cells, although far less abundant in tonsil cultures than lymphocytes, were edited with similar efficiency (macrophage CXCR4 surface levels: NT 3845.7±742.4 AU vs. KO 165.2±68.3; DC CXCR4 surface levels: 8464.0±640.9 vs. 67.4±21.1; supplementary Fig. S2F-J). Such highly efficient KO was consistently obtained with tonsil cultures from different donors, however the extent by which the procedure impaired cell viability varied between cells from different donors. Significant reduction in cell viability was accompanied by an upregulation of T cell activation markers CD25 and CD69, indicating general stress induced by the nucleofection (Fig. 3N-Q). Nevertheless, KO of CXCR4 in bulk tonsil cultures was functional and prevented productive HIV-1 infection, virus-induced CD4 T cell depletion and virus replication (Fig. 3R-X). Altogether, these results establish that genes can be efficiently knocked out in bulk tonsil cultures, however also reveal that this approach is associated with significant cytotoxicity and unspecific CD4 T cell activation.

**FIG 3.**
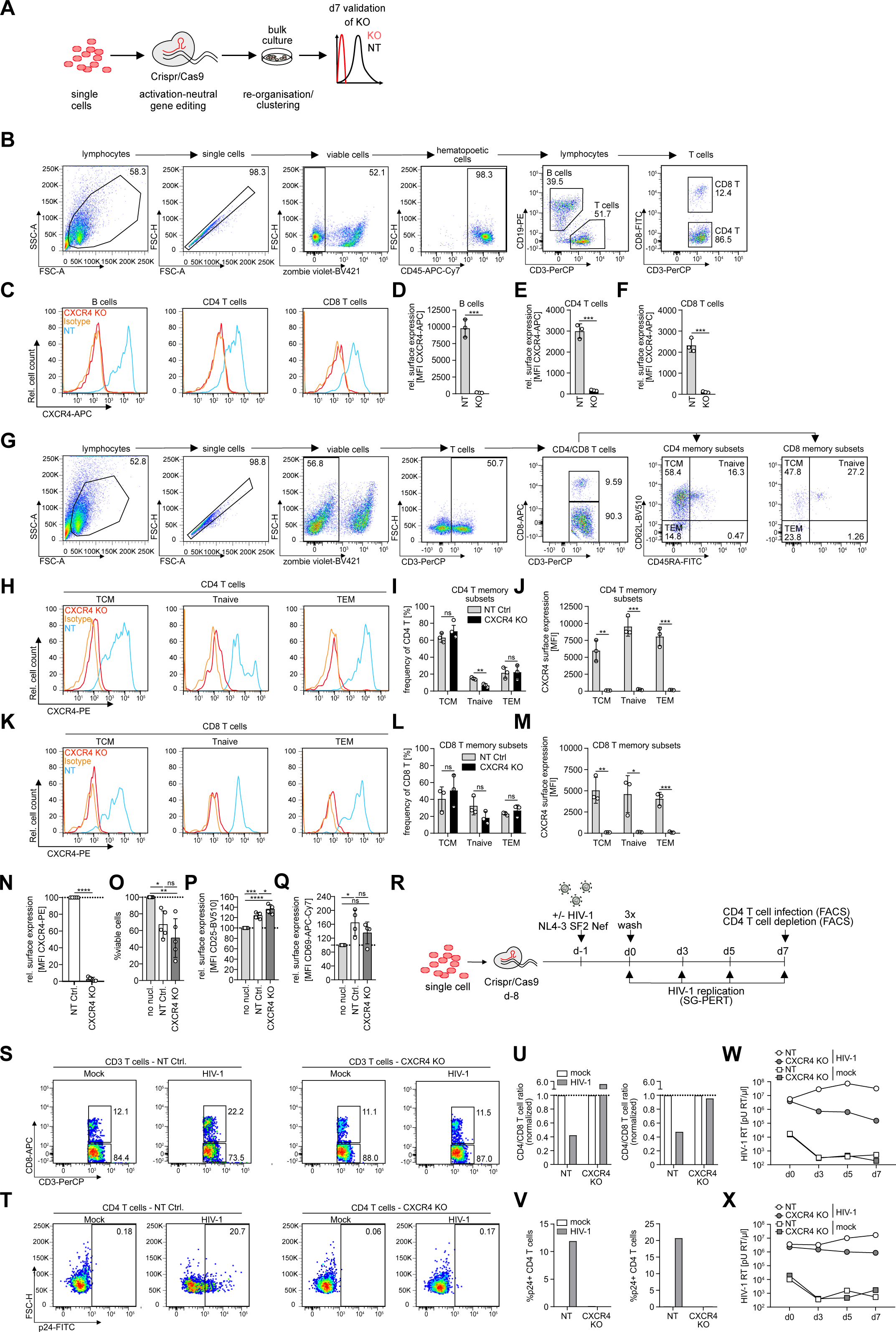
Gene editing of bulk tonsil cells by CRISPR/Cas9 nucleofection is highly efficient in all major tonsil cell populations and across T cell differentiation states. (A) Schematic of experimental setup. Tonsil cells were thawed, nucleofected with non-targeting or CXCR4 RNP complex and cultured in 96-well F-bottom plates for one week until analysis of KO efficiency in distinct cell populations by flow cytometry. (B) Representative dot plots and gating strategy for identification of lymphocyte subsets among tonsil cell cultures. (C) Representative histograms showing CXCR4 surface expression on B cells, CD4 and CD8 T cells on day 7 post nucleofection with CXCR4-targeting RNP (CXCR4 KO) with superimposed histograms for the non-targeting RNP (NT) condition and isotype control. (D)-(F) Quantification of C for NT or CXCR4 KO in B cells, CD4 or CD8 T cells. Each dot represents one tonsil. Shown are means with SD. (G) Representative dot plots and gating strategy for identification of memory subsets based on CD45RO and CD62L expression among CD4 and CD8 T cells in tonsil cell cultures. (H) and (K) Representative histograms showing CXCR4 surface expression on central memory (T_CM_), naïve (T_naive_) and effector memory (T_EM_) CD4 (H) or CD8 (K) T cell subsets on day 7 post nucleofection with CXCR4-targeting RNP (CXCR4 KO) with superimposed histograms for the non-targeting RNP (NT) condition and isotype control. (I) and (L) Distribution of memory subsets among CD4 (I) and CD8 (L) T cells for the NT or CXCR4 KO condition. (J) and (M) Quantification of H and K for NT or CXCR4 KO conditions in the respective memory subsets. Each dot represents one tonsil. Shown are means with SD. (N) Relative surface expression of CXCR4 on CXCR4 KO or NT tonsil cells on day 7 post nucleofection, calculated by normalizing the MFI in CXCR4 KO conditions to the NT control. (O)-(Q) Frequency of viable cells (O) or surface expression of CD25 (P) and CD69 (Q) on CD4 T cells among NT or CXCR4 KO cultures on day 7, normalized to non-nucleofected (no nucl.) cells, respectively. (N)-(Q) Each dot represents one tonsil. Shown are means with SD. Tonsils were cultured in 24-well transwell plates. (R) Schematic of experimental setup for analysis of HIV-1 infection in CXCR4 KO tonsil cultures. Briefly, tonsil cells were thawed, nucleofected with CRISPR/Cas9 RNP complex as indicated and infected with HIV-1 NL4.3SF2Nef at 1.5x10^5^ BCU per 2x10^6^ cells 7 days later by addition of the virus to the cultures. On the following day (d0), cells were washed 3x and infection was assessed by analysis of CD4 T cell infection and depletion by flow cytometry analysis upon final harvest and production of HIV-1 virions in the supernatant by SG-PERT throughout the infection course. (S) and (T) Representative dot plots showing CD4 and CD8 T cells subgated from CD3 T cells (S) as well as p24 signal in CD4 T cells (T) in NT or CXCR4 KO mock or HIV-1 infected conditions, respectively on day 7 post infection. (U) and (V) Quantification of S and T for two tonsils with the ratio of CD4 vs. CD8 T cells in U normalized to the mock condition for NT or CXCR4 KO, respectively. (W) and (X) HIV-1 RT activity in the supernatant of mock or HIV-1 infected NT or CXCR4 KO cultures at indicated timepoints for the two tonsils from S and T, respectively. Cells were cultured in 96-well V-bottom plates. Statistical significance was assessed by unpaired t-test for comparison of two groups or one-way ANOVA for comparison of more than two groups (ns, p>0.05; *, p<0.05; **, p<0.01; ***, p<0.001; ****, p<0.0001).

### Establishment of ediTONSIL gene KO in tonsil CD4 T cells

We reasoned that nucleofection conditions would need to be optimized for each cell population in tonsil cultures to limit cytotoxicity. Since there is no approach that renders nucleofection specific for a distinct cell subset in a complex heterogeneous mix of different cells, we attempted to improve cell viability following gene editing by isolating specific cell populations from tonsil cultures for editing and subsequent reuniting these cells with the unedited tonsil cells to reconstitute the original cell composition of the culture (Fig. 4A). Focusing on CD4 T cells, we first established positive selection of CD4 T cells by magnetic beads as the most efficient method for cell isolation (CD4 T cell purity 94.5/55.6% for two donors after negative isolation vs. 91.7/85.6% for the same tonsils after positive selection, respectively and 16.5/6.1% vs. 2.5/1.5% CD4 T cells left in the non-CD4 cell fraction; supplementary Fig. S3A-D) which enabled us to faithfully reconstitute tonsil cultures following isolation and add back (Fig. 4B). Importantly, efficiency of the positive isolation with respect to CD4 T cell depletion from the non-CD4 fraction is underestimated by negative gating for CD4 T cells based on CD3 and CD8 staining since a large fraction of the resulting population does not express CD4, likely representing unconventional T cells and further validating the efficiency of the CD4 positive selection (supplementary Fig. S3A-H). Of note, the positive isolation method did not affect viability, activation or susceptibility to HIV-1 infection (supplementary Fig. S3I-T). Nucleofection of isolated CD4 T cells with CXCR4-specific CRISPR/Cas9 RNP complexes and immediate reconstitution resulted in highly efficient reduction of CXCR4 on CD4 T cells within 7 days post nucleofection (mean of 22.4±10.1% CXCR4 surface expression left for KO normalized to NT CD4 T cells, Fig. 4C-D). Importantly, this procedure did not affect CD4 T cell viability or activation state and exogenous cytokine supplementation was thus not required to enhance survival, likely explained by the presence of T cell survival factors that are already provided by the tonsil microenvironment^36,37^ (Fig. 4E and supplementary Fig. S4A-F). Similar KO efficiency was obtained in positively vs. negatively isolated CD4 T cells, excluding adverse effects of the positive isolation strategy, e.g. by residual magnetic particles on CD4 T cells during nucleofection. Moreover, independently prepared RNP complexes yielded comparable KO efficiencies, validating the reproducibility of the workflow (supplementary Fig. S4G-I). Infecting tonsil cultures that were reconstituted after CXCR4 gene editing in CD4 T cells with HIV-1 (Fig. 4F) revealed that the removal of the entry co-receptor strongly reduced virus-induced depletion of CD4 T cells (∼5% relative CD4 T cell reduction in in CXCR4 KO vs. ∼75% in the NT condition) and fully abrogated virus replication in these cultures to a similar extend as treatment with the HIV-1 entry-inhibitor T20 (Fig. 4G-K and supplementary Fig. S4J-N).

**FIG 4.**
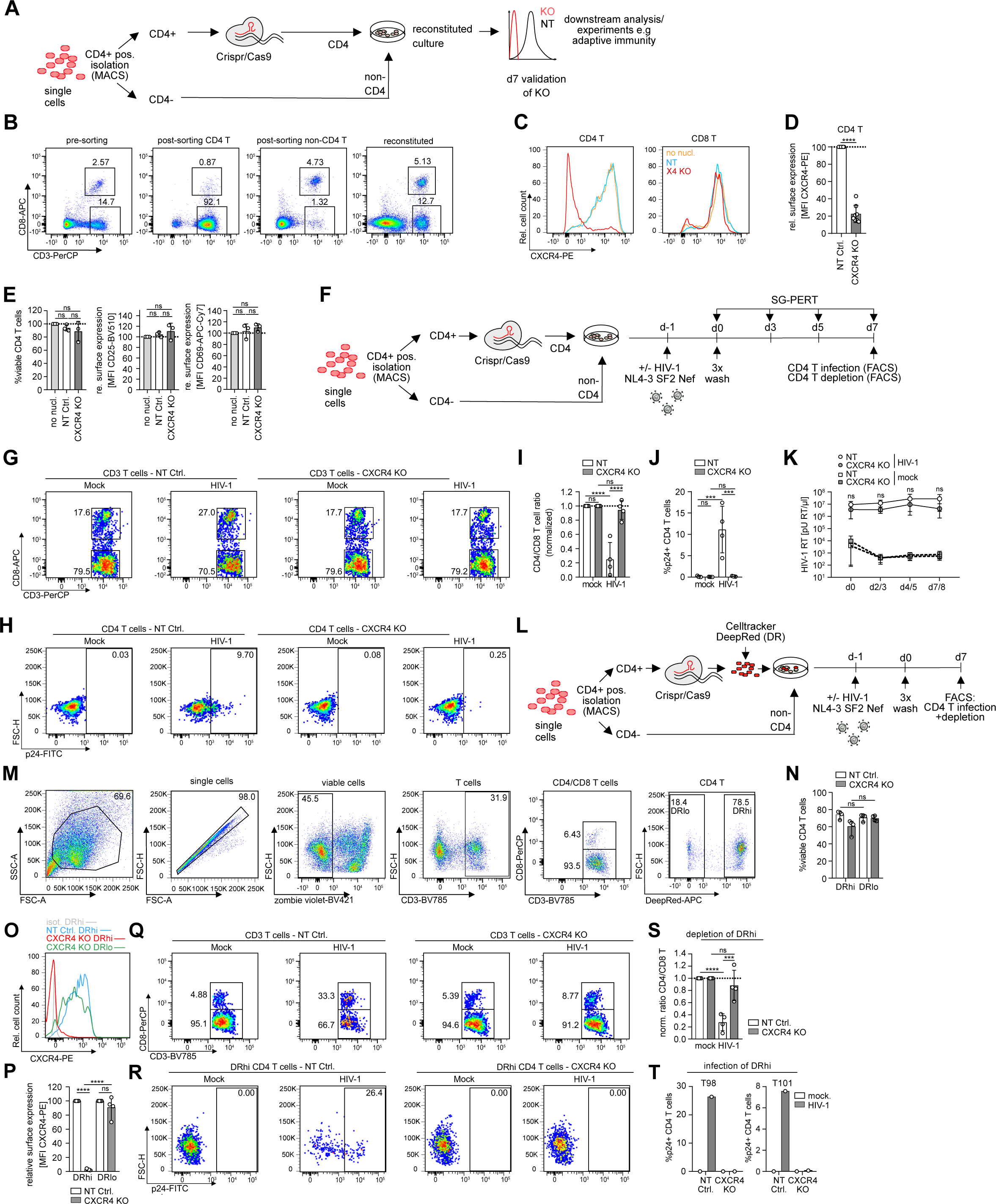
CD4 T cells can be specifically targeted for gene editing by CRISPR/Cas9 RNP nucleofection in tonsil culture by magnetic cell isolation and reconstitution. (A) Schematic of experimental setup. Briefly, tonsil cells were thawed and subjected to positive magnetic cell isolation of CD4 T cells. Magnetic beads were then removed from CD4 T cells prior to CRISPR/Cas9 RNP nucleofection and CD4 T cells were merged with the negative, non-CD4 T cell fraction, forming the reconstituted culture. 7 days post nucleofection, efficiency of the KO was assessed by flow cytometry. (B) Representative dot plots showing CD4 and CD8 T cell frequencies among bulk cells prior to CD4 sorting (pre-sorting) and CD4 fraction after sorting (post-sorting CD4 T) and non-CD4 fraction after sorting (post-sorting non-CD4 T) as well as the reconstituted culture post merging of CD4 and non-CD4 fraction (reconstituted). (C) Representative histograms of CXCR4 expression on CD4 and CD8 T cells on day 7 following CD4 T cell isolation, nucleofection and merging with non-CD4 T cells. Shown is the CXCR4 surface signal on non-nucleofected (no nucl.), NT control (NT) or CXCR4 KO (X4 KO) cultures. (D) Quantification of CXCR4 on CD4 T cells for 8 tonsils. CXCR4 signal was normalized to the NT condition for each tonsil. Tonsil cells were cultured in 24-well transwell or 96-well V-bottom plates. (E) Frequency of viable CD4 T cells and surface expression of CD25 and CD69 among NT or CXCR4 KO CD4 T cells on day 7 for 3 tonsils, normalized to non-nucleofected cells, respectively. Tonsil cells were cultured in 24-well transwell plates. (D), (E) Each dot represents one donor. Shown are means with SD. (F) Schematic of experimental setup to assess HIV-1 infection in tonsil cultures with edited CD4 T cells. Briefly, tonsil cells were thawed and CD4 T cells were isolated and nucleofected prior to merging with non-CD4 T cells. On day 7 post nucleofection, cultures were infected with HIV-1 NL4.3SF2Nef at 1.5x10^5^ BCU per 2x10^6^ cells and HIV-1 infection was assessed on subsequent days by analysis of CD4 T cell depletion, CD4 T cell infection by intracellular p24 staining and production of new virions in the supernatant by SG-PERT. (G) and (H) Representative dot plots showing frequencies of CD4 and CD8 T cells among tonsil CD3 T cells (G) or intracellular p24 staining among tonsil CD4 T cells (H) in mock or HIV-1 infected cultures on day 7 post infection for NT or CXCR4 KO cultures, respectively. (I) Quantification of G for 4 tonsils, normalized to the mock condition, respectively. (J) Quantification of H for 4 tonsils. Tonsil cells were cultured in 24-well transwell or 96-well V-bottom plates. (I), (J) Each dot represents one donor. Shown are means with SD. (K) SG-PERT quantification of HIV-1 RT activity in the supernatant at indicated timepoints for mock or HIV-1 infected cultures nucleofected with NT or CXCR4 KO RNP. Shown are means with range of 3 donors per condition. Tonsil cells were cultured in 24-well transwell or 96-well V-bottom plates. (L) Schematic of the experimental set-up for identification of nucleofected CD4 T cells among total CD4 T cells in tonsil culture by celltracker staining. (M) Representative dot plots and gating strategy for identification of nucleofected, celltracker Deepred high intensity (DRhi) CD4 T cells in tonsil bulk cultures. (N) Quantification of viable CD4 T cells among DRhi and DRlo cells in NT and CXCR4 KO conditions for 4 tonsils. (O) Representative histogram of CXCR4 expression on celltracker-stained (DRhi), CXCR4 KO CD4 T cells with superimposed histogram of the DRhi population in the NT condition and isotype control as well as DRlo CD4 T cell population in the CXCR4 KO condition. (P) Relative surface expression of CXCR4 on the DRhi and DRlo population in CXCR4 KO tonsil CD4 T cells on day 7 post nucleofection, normalized to the NT control for 4 tonsils. Shown are means with SD. (Q) and (R) Representative dot plots showing frequencies of CD4 and CD8 T cells among tonsil CD3 T cells (Q) or intracellular p24 staining among DRhi tonsil CD4 T cells (R) in mock or HIV-1 infected cultures on day 7 post infection for NT or CXCR4 KO cultures, respectively. (S) Quantification of the CD4 vs. CD8 T cell ratio as calculated for the DRhi population among total CD4 T cells for 4 tonsils, normalized to the mock condition, respectively. (N),(P),(S) Each dot represents one donor. Shown are means with SD. (T) Quantification of R in HIV-1 infected cultures for 2 tonsils as depicted. Tonsil cells for celltracker experiments were cultured in 96-well V-bottom plates. Statistical significance was assessed by unpaired t-test for comparison of two groups or one-way ANOVA for comparison of more than two groups (ns, p>0.05; *, p<0.05; **, p<0.01; ***, p<0.001; ****, p<0.0001).

The KO procedure was highly efficient as only ∼1% of CD4 T cells in the reconstituted cultures was precluded from editing. These represent residual CD4 T cells present in the non-CD4 T cell fraction that could not be removed during initial cell separation (Fig. 4B and supplementary Fig. S3E-H). While not sufficient to drive appreciable HIV-1 replication in CXCR4 KO cultures, this residual population may exert immune function that are relevant for other scenarios. We therefore tested if labeling of the gene-edited CD4 T cell population allows to phenotypically distinguish edited and unedited cell population in reconstituted tonsil cultures (Fig. 4L). Staining of gene-edited CD4 T cells prior to culture reconstitution with DeepRed celltracker followed by one week of culture indeed allowed the identification of gene edited (DeepRed high (DRhi) and unedited (DeepRed low (DRlo) populations with unaffected viability of celltracker-stained cells (74.2±4.4% viable cells in NT DRhi CD4 T vs. 71.1±3.9% in DRlo and 60.2±8.3 % in CXCR4 KO DRhi CD4 T vs. 69.7±3.2% in DRlo; Fig. 4M, N). Of note, CXCR4 cell surface expression was undetectable on DRhi cells (Fig. 4O, P), revealing that the residual CXCR4^+^ CD4 T cell population in the experiment described above (Fig. 4D) reflects the presence of unedited cells, largely consisting of CD4/CD8-negative T cells and a minor fraction of CD4 T cells that resisted positive selection. Consistently, DRhi cells were fully protected from HIV-1 infection (Fig. 4Q-T). Isolation of CD4 T cells and subsequent reconstitution of tonsil cultures thus allows for highly efficient and activation neutral gene editing where edited cell populations can be further labeled by celltracker staining to unambiguously identify cells of interest while enabling direct comparison with unedited cells in the same condition.

### Gene editing to study complex adaptive immune reactions

In contrast to HIV-1 replication and virus-induced depletion of CD4 T cells, complex immune reactions such as mounting of recall CD8 T cell responses (supplementary Fig. S1F-P) depend on functional interactions of several immune cell types. We therefore assessed next the impact of ediTONSIL on adaptive immune reactions. First, we performed RNP nucleofection on bulk tonsil cultures or on isolated CD4 T cells with subsequent culture reconstitution and assessed the response of these cultures to T cell activation by SEB (Fig. 5A). Under both conditions, the analysis of cell surface activation markers revealed an efficient response of CD4 T cells to stimulation that was not significantly altered in magnitude and kinetic by the nucleofection procedure (Fig. 5B-E). We next applied the analogous experimental setting to the expansion of EBV-specific CD8 T cells in response to the EBV peptide pool (Fig. 2G, 5F). While RNP nucleofection of bulk tonsil cells abrogated the ability of these cultures to expand IFNγ+ CD8 T cells, editing of isolated CD4 T cells did not impair their ability to provide help for CD8 T cell expansion (Fig. 5G-I). Consistently, the CD8 T cell subset distribution was skewed towards effector cells (T_Eff_) upon bulk nucleofection while isolated editing of CD4 T cells did not significantly affect the memory T cell subset composition of these cultures (Fig. 5J-O). EdiTONSIL of isolated tonsil CD4 T cells thus preserves the capacity of reconstituted HLACs to undergo complex adaptive immune reactions.

**FIG 5.**
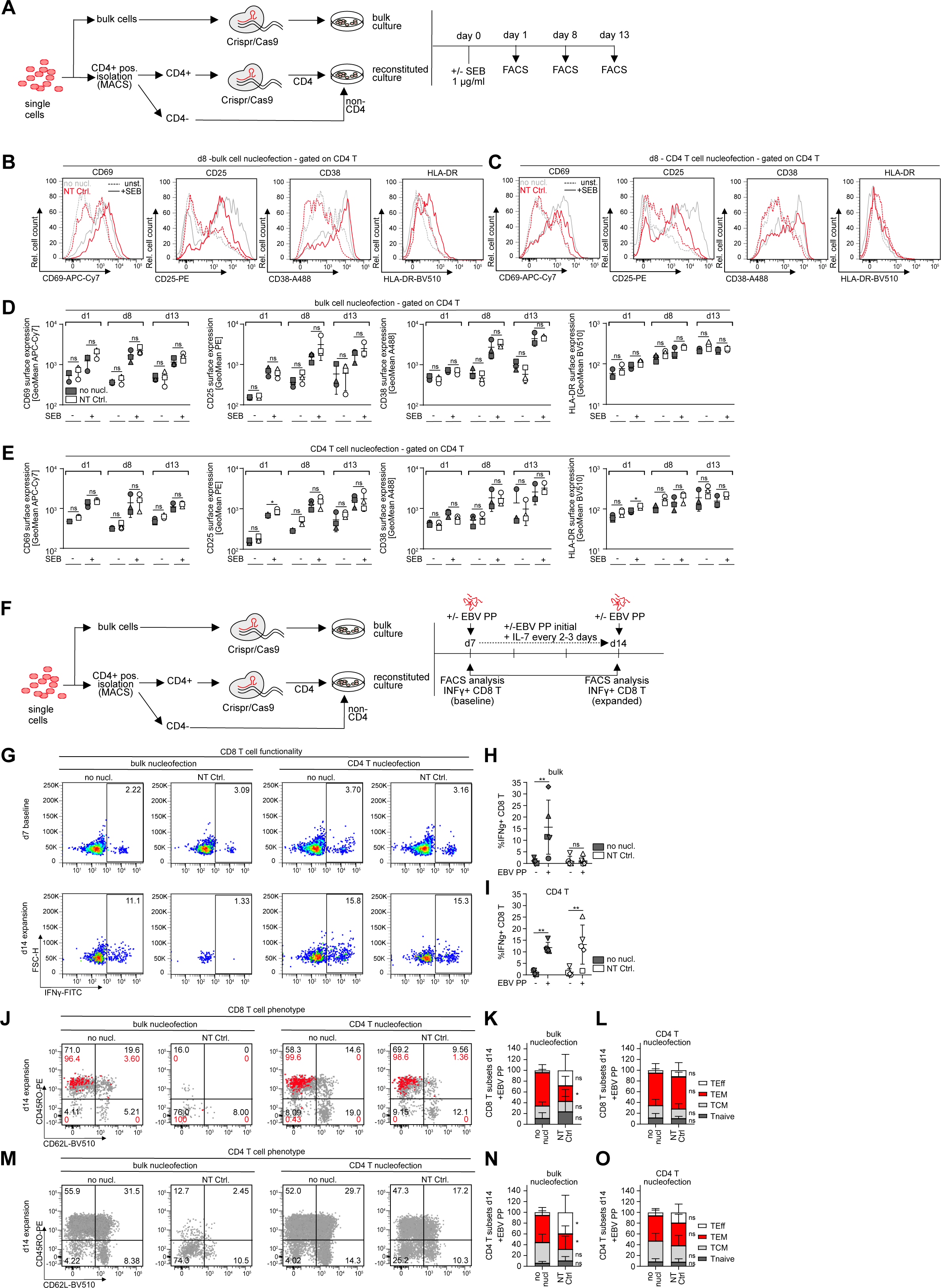
CRISPR/Cas9-mediated gene editing of tonsil CD4 T cells can be used as a tool to mechanistically dissect T cell activation and CD4 T cell helper function. (A) Schematic of the experimental set-up to assess activation response of CD4 T cells after nucleofection of bulk or isolated CD4 T cells. Briefly, bulk cells or positively isolated CD4 T cells after bead removal were either left untreated or nucleofected with NT RNP complex prior to seeding, i.e. merging with non-CD4 T cells for the reconstituted culture. One day after seeding, cultures were stimulated with SEB or left untreated and activation response was assessed on subsequent days by flow cytometry. (B) and (C) Representative histograms showing CD69, CD25, CD38 and HLA-DR expression on tonsil CD4 T cells on day 7 post nucleofection of bulk cells (B) or isolated CD4 T cells (C). Superimposed histograms show expression profiles of the non-nucleofected culture (grey) and cultures nucleofected with non-targeting control (NT Ctrl) RNP. Dashed lines represent unstimulated conditions while solid lines represent SEB-stimulated cultures. (D) and (E) Quantification of B and C for day 1, day 8 and day 13 of culture for non-nucleofected or NT conditions with and without SEB stimulation after bulk (D) or isolated CD4 T cell (E) nucleofection. Each symbol represents one donor. Shown are means with SD. Tonsil cells were cultured in 96-well F-bottom plates. (F) Schematic of the experimental set-up for assessing immunocompetency of tonsil bulk or CD4 T cells after nucleofection. Briefly, bulk cells or positive-isolated CD4 T cells after bead removal were either left untreated or nucleofected with NT RNP complex prior to seeding, i.e. merging with non-CD4 T cells for the reconstituted culture. Seven days after seeding, EBV peptide pool was added together with IL-7. On day 7 and day 14, baseline and expanded EBV-specific CD8 T cells, respectively, were identified by production of IFNγ following stimulation with the EBV peptide pool. (G) Representative dot plots showing IFNγ+ CD8 T cells in tonsil cultures for bulk or isolated CD4 T cell nucleofection (NT Ctrl.) in comparison to untreated (no nucl.) conditions on day 7 or day 14 of culture upon re-stimulation with EBV PP, respectively. (H) and (I) Quantification of (G) for five tonsils, showing frequency of IFNγ+ CD8 T cells on day 14 in cultures were either bulk cells or isolated CD4 T cells were left untreated (no nucl) or nucleofected with NT Ctrl. Shown are means with SD. Each symbol represents one donor. (J) and (M) Characterization of memory subsets based on expression of CD45RO and CD62L among total CD8 (J) or CD4 (M) T cells (grey) or superimposed IFNγ+ CD8 T cells (J)(red) on day 14 following peptide stimulation of tonsil cells cultured with EBV peptide pool for 7 days. (K) and (L) Quantification of (J) for total CD8 T cells in cultures where bulk cells (left panel) or isolated CD4 T cells (right panel) were nucleofected (NT Ctrl.) or left untreated (no nucl.). (N) and (O) Quantification of (M) as described above. T_Eff_ = T effector cells, T_EM_ = effector memory cells, T_CM_ = central memory cells, T_naive_ = naïve T cells. Shown are means with SD for five tonsils. Tonsil cells were cultured in 96-well F-bottom plates. Statistical significance was assessed by one-way ANOVA (ns, p>0.05; *, p<0.05; **, p<0.01; ***, p<0.001; ****, p<0.0001).

## Discussion

Increasing efforts are made to study pathobiological principles and their therapeutic targeting in complex organoid and organotypic models to account for the impact of tissue environments and cell type heterogeneity at complexity levels between individual cell types (cell line or isolated primary cells) and small animal models / patient samples. In particular for lymphoid organs that predominately consist of hematopoietic cells, which are difficult to generate by iPS technology^38,39^, *ex vivo* tonsil cultures are a well-established organotypic model that proved invaluable to study complex pathogen replication and complex immune reactions *ex vivo*^5,7,9,18–20,24,40–43^. However, the full potential of this culture model for gaining molecular insight into immune reactions could so far not be exploited due to the lack of gene editing methods that allow for modification of gene expression with high precision and without impairing the activation state and immune competence of tonsil-resident cells. In the case of HLAC, a particular challenge for such an approach lies in the fact that close to 100% KO efficiency is required to conduct meaningful studies, since even minor contamination with unedited subpopulations may be sufficient to exert complex functions, expand in the course of the experiment and thereby potentially mislead conclusion on the relevance of the gene of interest in the process studied. Furthermore, the gene editing approach per se should not affect the intrinsic properties of the edited cells. The ediTONSIL workflow established herein offers these possibilities as it allows gene editing with over 98% efficacy without requiring exogenous activation. Editing can in principle be done in bulk cultures to effectively target all major cell populations in parallel, although this comes at the cost of a significantly reduced cell viability. However, due the fact that HLACs are launched as suspension cultures that subsequently reorganize into immune competent structures, specific cell populations can be isolated and edited at the beginning of an experiment. Importantly, ediTONSIL of isolated CD4 T cells neither causes cell activation per se nor does it impair the activation response to external stimulation such as antigenic stimulation of CD4 T cells and also preserves the competence for complex functional cell-cell interaction such as CD4 T cell help to a recall CD8 T cell response. As illustrated at the example of HIV-1 infection, kinetics and magnitude of virus replication as well as CD4 T cell depletion are unaffected by the editing approach but targeting essential host cell genes such as the entry co-receptor fully abrogates virus infection and spread. Adapting the methodology established here for CD4 T cells to e.g. CD8 T cells, B cells and myeloid cells will offer a versatile toolbox to study mechanisms in all cell types present in the culture. This, for the first time, allows precise dissection of molecular mechanisms in basic immunology but also complex cell-cell interactions in the tissue context. This may instrumental for e.g. for dissecting the mechanisms underlying CD4 T cell help to CD8 T cell responses, which is an emerging concept in infectious diseases and cancer^44–47^ and insights yielded from ediTONSIL studies in a human lymphoid tissue system will complement *in vivo* mouse studies. Importantly, this culture system is also suitable to study humoral immunity, e.g. B cell antibody production to recall antigens but also *de novo* vaccine responses and ediTONSIL opens new possibilities to study B cell responses^7,19,20^. Furthermore, studies on viral pathogenesis and cell interactions in *ex vivo* tonsil cultures, exemplified herein with HIV-1, can be expanded to other pathogen infections including HTLV, human Herpesviruses (HHV-6, HHV-7, HCMV, HSV-2, EBV), vaccinia virus, measles virus and West-Nile virus^17,40–43,48–53^. ediTONSIL thus opens new avenues towards the mechanistic dissection in organotypic lymphoid cultures and will be widely applied to studies of pathogen-host interaction but also of fundamental principles of adaptive immunity.

### Limitations of the study

The workflow developed herein allows for the efficient gene editing of bulk or individual cell populations in *ex vivo* tonsil cultures. For individual cell populations however, the isolation efficiency is a critical parameter and the success of ediTONSIL depends on the availability and quality of commercial or in-house isolation procedures for cells of interest. Furthermore, it is important to note that the half-life of each individual target protein is a key determinant. Since the viability of these *ex vivo* cultures significantly decreases after two weeks, proteins with low turnover will not be sufficiently depleted within the live span of the culture. The fact that the applicability of this workflow depends of these individual features of each target implies that KOs have to be monitored specifically for each target, which currently precludes the use of this method for high throughput screening approaches. Establishing conditions that expand the life span of these model cultures will be a central goal of future developments.

## Acknowledgments

This project is supported by the Deutsche Forschungsgemeinschaft (DFG, German Research Foundation), Projektnummer 240245660 - SFB 1129 (project 8) and Projektnummer 259021520 – FA 378/13-2, OK 742/5-2) and the German Centre for Infection Research (DZIF) (TTU 04.820 – HIV reservoir to OTF, OTK. OTF is a member of the CellNetworks cluster of excellence (EXC81). We want to especially thank Nadine Tibroni, Ina Ambiel and the Head and Neck Surgery team for support with the acquisition of tonsil specimens and all anonymous tonsil donors that made this study possible.

## Author Contributions

Conceptualization, O.T.F.; Methodology, L.S., and O.T.K.; Investigation, K.M.; Visualization: K.M., Writing – Original Draft, O.T.F. and K.M. Writing –Review & Editing, O.T.F, K.M., G.D. and O.T.K; Funding Acquisition, O.T.F., O.T.K.; Resources, D.G.; Supervision, O.T.F.; Project administration: O.T.F.

## STAR Methods

### RESOURCE AVAILABILITY

#### Lead Contact

Further information and requests for resources and reagents should be directed to and will be fulfilled by the Lead Contact, Prof. Dr. Oliver T. Fackler.

#### Materials Availability

There are no new materials generated in this paper. All material has to be requested from authors of cited references.

#### Data and Code Availability

Primary data from all FACS analysis generated in this study are available from the Lead Contact upon request.

### EXPERIMENTAL MODEL AND SUBJECT DETAILS

#### Human lymphoid aggregate culture (HLAC)

HLACs were generated as previously described^19^. Tonsils were removed and received from randomly selected anonymous donors with informed consent in accordance with the ethics vote S-123/2014 from the Heidelberg University Hospital ethics committee. Gender and age of the donors is not known. One half of each tonsil was collected during standard tonsillectomy while the other half was processed in the department of pathology to assess chronic tonsillitis and exclude any unexpected malignancy. Removed tonsil tissue was immediately transferred to 50ml Falcon tubes containing 1X phosphate-buffered saline (PBS) and processed on the same day as described^19^. In brief, dead and burnt tissue was removed and tonsils were sliced into blocks of approximately 2mm size that were passed through a 100 µm mash PET strainer. The suspension was filled up with PBS to 35 ml and erythrocytes and debris was removed from the cell suspension by layering onto 15 ml of Biocoll Separating Solution and centrifugation for 20 min. at 2000 rpm (Heraeus MEGAFUGE 40R) without break at room temperature. The mononuclear fraction above the erythrocyte layer was isolated, washed twice with PBS and frozen in 90% FBS with 10% DMSO at -80°C in aliquots of 1x10^8^ cells/ml. For experiments, cells were thawed in tonsil medium (RPMI 1640 GlutaMAX, 10% FCS, 5% Fungizone, 5% non-essential amino acids, 5% Sodium Pyruvate, 5% Penicillin/Streptomycin, 5% ITS-G), washed twice and resuspended in 10 ml medium prior to viability and cell counting in a 1:100 dilution with Trypan blue. Cells were resuspended in tonsil medium at 1x10^7^ cells/ml and 200 µl of the cell suspension was added per well of a V or F bottom 96-well plate. For transwell culture in 24-well plates, cells were resuspended at 4x10^7^ cells/ml and 50µl cell suspension was added to the upper transwell while the lower well was filled with 500 µl medium. Transwells were equilibrated with medium prior to use.

Cells were cultured at 37°C, 5% CO_2_ under humidified conditions for up to three weeks.

#### Cell culture

Human embryonic kidney 293T (HEK293T) cells were cultured in Dulbecco’s modified Eagle medium (DMEM; ThermoFisher Scientific) containing 10% FBS, 50 U/ml penicillin and 50 μg/ml streptomycin. The cells are female and have not been authenticated. TZM-bl cells (JC53BL-13) were obtained from the NIH AIDS Research and Reference Reagent Program (Cat. no. 8129) and are derived from a HeLa cell clone that was engineered to express CD4, CCR5 and CXCR4 to allow entry of X4- and R5-tropic HIV-1 strains and further contains integrated reporter genes for firefly Luc and *E. coli* β-galactosidase under the control of an HIV-1 long terminal repeat. For determination of virus stock infectivity, TZMbl cells were infected with titrated amounts of the virus and X-β-Gal substrate was added after 2-3 days. The substrate is converted by the β-galactosidase to a blue colored product that is used to assess infection of TZMbl cells and relative infectivity of the virus stock.

### METHOD DETAILS

#### Virus production, infectivity measurements, infection assays and CD4 T cell depletion

Virus production from 293T cells, infectivity measurements using SG-PERT for total viral particles and TZM-bl luciferase reporter assay were performed as described previously^27,28^. 2x10^6^ tonsil cells were infected with 1.5x10^5^ BCU/well of the proviral CXCR4 tropic chimera HIV-1_NL4.3_ SF2 Nef WT (X4 HIV-1). For F- and V-bottom 96-well plates, virus was diluted in 100 µl/well and added after removal of 100 µl spent medium from each well. For transwell plates, 250µl medium were removed from the lower transwell and virus was added in 250µl fresh medium. Mock wells were treated accordingly without addition of virus to the medium. One day post-infection (dpi) HLACs were washed 3x with 200 µl tonsil medium for V- and F-bottom 96-well plates by centrifugation at 1200 rpm for 5 min. at RT and addition of fresh medium after washing. Transwell cultures were washed by 3x renewal of the lower transwell medium (500 µl) with 1 min. incubation time respectively to allow full submerging of the upper transwell and transition of input virus in the supernatant to the lower well. HLACs were cultured for 7-15 days post infection. Every two to three days, 5 µl supernatant for determination of viral titers was harvested. The measurement started 1 dpi with the initial supernatant collected after the washing step.

#### Flow cytometry analysis

Tonsil cells were washed with PBS and stained with zombie violet fixable viability dye (1:1000 in PBS), incubated for 15 min. at RT and washed with FACS buffer (1xPBS +0.5% BSA +2mM EDTA) prior to surface antibody staining. Surface staining of CD45, CD3, CD4, CD8, CD19, HLA-DR, CD11b, CD11c, CD38, CD25, CD69, CD45RO, CD45RA, CD62L and CXCR4 was performed at a 1:100 dilution of antibodies in FACS buffer and incubation at 4°C for 20 min, respectively. Cells were then washed and fixed with 3% PFA for 10 min. or 90 min. for BSL-3 sample inactivation. For intracellular staining of HIV-1 capsid (p24), cells were then resuspended in 0.1% Triton X-100 containing 1:100 diluted anti-p24 antibody (clone KC57, Becton Coulter) and incubated for 20 min. at 4°C and acquired on the same day. For intracellular staining of IFNγ, cells after surface staining and PFA fixation were permeabilized and stained using the BD Cytofix/Cytoperm kit according to the manufacturer’s protocol. Briefly, cells were incubated in Cytofix/Cytoperm solution for 20min. at 4°, washed 2x with 1xPerm buffer and resuspended in 1xPerm buffer containing 1:100 diluted anti-human IFNγ antibody. Samples were incubated for 30 min. at 4°C, washed and acquired by flow cytometry on the same day.

Samples were acquired using a BD FACSVerse with BD FACSuite Software or FACS Celesta with BD FACS Diva software. Gating was performed using FlowJo software 10.4.2 and data was processed with Microsoft Office Excel 2016 and GraphPad Prism 6.0 software.

#### Quantification of HIV-1 induced CD4 T cell depletion

CD4 T cell depletion was quantified as described previously^10^. In brief, the CD4/CD8 ratio was calculated for each sample and related to the ratio of the corresponding uninfected mock sample that was set to 1.

#### RNP preparation

RNP complexes were prepared as described^21^. Briefly, sgRNAs were obtained from Synthego and reconstituted to 100 µM in TE buffer supplied by the manufacturer. Cas9 was obtained from IDT in a ready-to-use solution of 62 µM. SgRNA and Cas9 were mixed on ice at a ratio of 2.5:1 (sgRNA:Cas9) with PBS to obtain final concentrations of 100 pmol of sgRNA and 40 pmol of Cas9 in 5 µl complex volume, incubated for 15-20 min. at RT and frozen at -80°C until usage. Immediately before nucleofection, the RNP complexes for NT Ctrl. or CXCR4 were thawed on ice. Equal amounts of NT Ctrl. RNP and CXCR4 RNP were used per cell in each experiment for proper comparison. The final CXCR4 RNP mix with which cells were nucleofected further consisted of equal amounts of two different sgRNA-Cas9 RNP complexes as described^21^.

#### CRISPR/Cas9 RNP nucleofection of tonsil cells

Tonsil bulk cells or isolated and bead-released CD4 T cells were prepared for nucleofection as described (ref. Albanese et al. 2021). Briefly, cells were washed once in 1xPBS and resuspended in P3 primary cell nucleofection buffer (Lonza) as recommended by the manufacturer’s protocol, using 2x10^6^ cells in 20µl nucleofection buffer for 20 µl Nucleocuvette Strip or 1x10^7^ cells in 100µl for single Nucleocuvettes. Cells were mixed briefly with the RNP mix that was prepared shortly before nucleofection on ice and were transferred to the respective nucleofection cuvettes. For 2x10^6^ cells, 6 µl RNP mix in total were used while 24µl were used for 1x10^7^ cells. Lonza 4D-Nucleofector® X Unit was used for electroporation and cells were immediately after resuspended in unsupplemented RPMI 1640 to reach a concentration of 4x10^7^ cells/ml and incubated for 15 min. prior to dilution in tonsil medium to a final concentration of 1x10^7^ cells/ml prior to merging with non-CD4 T cells. After nucleofection, cells were cultured either in 24-well transwell or 96-well V-/F-bottom plates as indicated. Culture plate set-up did not have any influence on KO efficiency.

#### SEB stimulation of tonsil cell cultures

SEB was added to tonsil cultures at 1µg/ml final concentration directly into the medium of V or F bottom 96-well plates or into the lower transwell of transwell plates and was left on the cells for the remaining culture period.

#### Cytokine treatment of tonsil cell cultures for optimization of post-nucleofection conditions

After nucleofection of CD4 T cells, reconstituted cell suspensions were seeded in 24-well transwell plates with IL-7, IL-15 and IL-2 alone or in combination (IL-7+IL-15) at indicated concentrations of 0.5 or 2ng/ml for IL-7 and IL-15 and 10 or 50 IU/ml for IL-2. Every 2-3 days, medium was refreshed by preparing a 3x concentrated pre-dilution of the concentrated cytokine stocks in tonsil medium which was added to fill up the lower transwells after removal of a third of the spent medium every 2-3 days.

#### T20 treatment of tonsil cells to inhibit fusion and entry of HIV-1

T20 (Fuzeon, Enfuvirtide) was added to the diluted virus stock for infection of tonsil cultures at 20 µM final concentration on day 7 post nucleofection, was added freshly after washing of the virus input and was renewed on day 4 post infection.

#### CD4 positive and negative isolation from tonsil cell suspension

In the experiment directly comparing the sorting and nucleofection efficacy in CD4 negative vs. positive sorted CD4 T cells (Fig. S3), tonsil cell suspensions were processed for positive or negative CD4 T cell isolation using stemcell EasySep™ Human CD4+ T Cell Isolation Kit or EasySep™ Release Human CD4 Positive Selection Kit according to the manufacturer’s recommendations. Briefly, tonsil cells were thawed, counted and concentration adjusted to 5x10^7^ cells/ml for negative isolation, i.e. 1x10^8^ cells/ml for positive isolation in 1xPBS with 2% FBS and 2 mM EDTA. All steps were followed according to the manufacturer’s protocol, yielding an unlabeled CD4 T cell fraction from the negative isolation and a CD4 antibody and magnetic-bead labeled CD4 T cell fraction from positive isolation which underwent bead removal with the supplied release buffer directly after isolation. Purity and efficiency of the isolation procedures were assessed using CD3 and CD4 antibody staining for the negative isolation and CD3 and CD8 staining for the positive isolation as recommended. Unlabeled non-CD4 T cells resulting from the CD4 positive isolation procedure (negative fraction) were used to complement both negative and positive isolated CD4 T cells after nucleofection for the reconstituted culture in Fig. S3 D to avoid a bias resulting from the presence of magnetic beads and possible induction of activation in cultures with negative isolated CD4 T cells and magnetic bead-labeled non-CD4 T cells. Non-CD4 T cells resulting from the positive CD4 T cell isolation were resuspended in tonsil medium, counted and resuspended at 1x10^7^ cells/ml before merging with CD4 T cells for downstream experiments.

For all experiments except the direct comparison of positive vs. negative CD4 T cell isolation, the non-CD4 T cell fraction underwent a second positive isolation procedure to deplete remaining CD4 T cells from this population or alternatively, double amounts of CD4 isolation cocktail and magnetic particles were used in the first positive isolation process in order to increase purity of the non-CD4 fraction which routinely decreased the frequency of contaminating CD4 T cells in this fraction to 1/5^th^ to 1/6^th^ of the original CD4 T cell population. After isolation, CD4 T cells were resuspended in tonsil medium, counted and processed for downstream applications, e.g nucleofection before resuspension at 1x10^7^ cells/ml and merging with unlabeled non-CD4 T cells at the same concentration, yielding a typical ratio of 1 to 4 (20%) CD4 T cells to non-CD4 T cells in the final reconstituted culture.

#### Celltracker staining of isolated tonsil CD4 T cells

Directly after nucleofection and incubation in unsupplemented RPMI 1640 at 4x10^7^ cells/ml, tonsil CD4 T cells were resuspended at 1x10^7^ cells/ml in celltracker DeepRed (Thermo Fisher) solution by addition of 3x volume of PBS containing 4x concentrated celltracker dye, were incubated for 15 min. at 37°C and were washed in tonsil medium 2x before resuspension at 1x10^7^ cells/ml and merging with unlabeled, non-nucleofected non-CD4 T cells for reconstituted cultures.

#### Reactivation and expansion of EBV-specific memory CD8 T cells in tonsil tissue

Tonsils used for CD8 T cell reactivation experiments were pre-selected in separate screening experiments based on the presence of EBV-specific CD8 T cells measured by EBV peptide-specific IFNγ production compared to background activation induced by unspecific HIV-1 Gag peptides. The EBV peptide pool (EBV Peptivator, Miltenyi Biotec) contains MHC class I and class II-restricted epitopes from EBV lytic and latent cycle proteins for stimulation of CD4 and CD8 T cells, respectively. The peptides contained in the pool are recognized by most common HLA types such as HLA-A*A02:01 and HLA-B*07:02, ensuring broad response rates across different tonsil donors. The peptide mix was directly added to the culture medium at 1 µg/ml (0.6 µM for each peptide) at indicated timepoints without disturbing the cell layer and IL-7 was added at a final concentration of 1 ng/ml. Cultures were incubated for 7 days with the peptide pool and IL-7 was refreshed every 2-3 days by removing a third of the spent medium and replacing with fresh medium containing 3x concentrated IL-7. To assess IFNγ-production as a correlate for EBV-spec. CD8 T cell reactivation at designated timepoints, separate wells were exposed to the peptide pool on the start day of culture (baseline frequency of reactive CD8 T cells) and after 7 days for 30 min. and subsequently incubated with Brefeldin A at 5 µg/ml final concentration for an additional 2 h to block cytokine export. Cells were then harvested with PBS +2 mM EDTA, washed with PBS and stained with zombie violet fixable viability dye and surface antibodies for human CD3, CD8, CD45RO and CD62L prior to fixation, permeabilization and intracellular cytokine staining. In every experiment, control wells that did not receive the peptide pool but were also stimulated at the end day of culture with the EBV peptide pool served as background controls for unspecific activation and expansion of CD8 T cells over the culture period that was negligible in our experiments. Further, antigen-specificity of the IFNγ production was validated in several experiments by stimulating the EBV-peptide pool cultures with an MHC class I epitope pool derived from HIV-1 Gag peptides in parallel, to which the healthy donors in our experiments did not respond with IFNγ production (data not shown).

#### Quantification and statistical analysis

Statistical analysis of datasets was carried out using Prism version 8.0.1 (GraphPad). Statistical significance was calculated using One-way ANOVA for comparison of more than one condition or unpaired t-test for comparison of two conditions as indicated. n.s., not significant; *, p < 0.05; **, p < 0.01; ****, p<0.0001. See figure legends for details.

### KEY RESOURCE TABLE

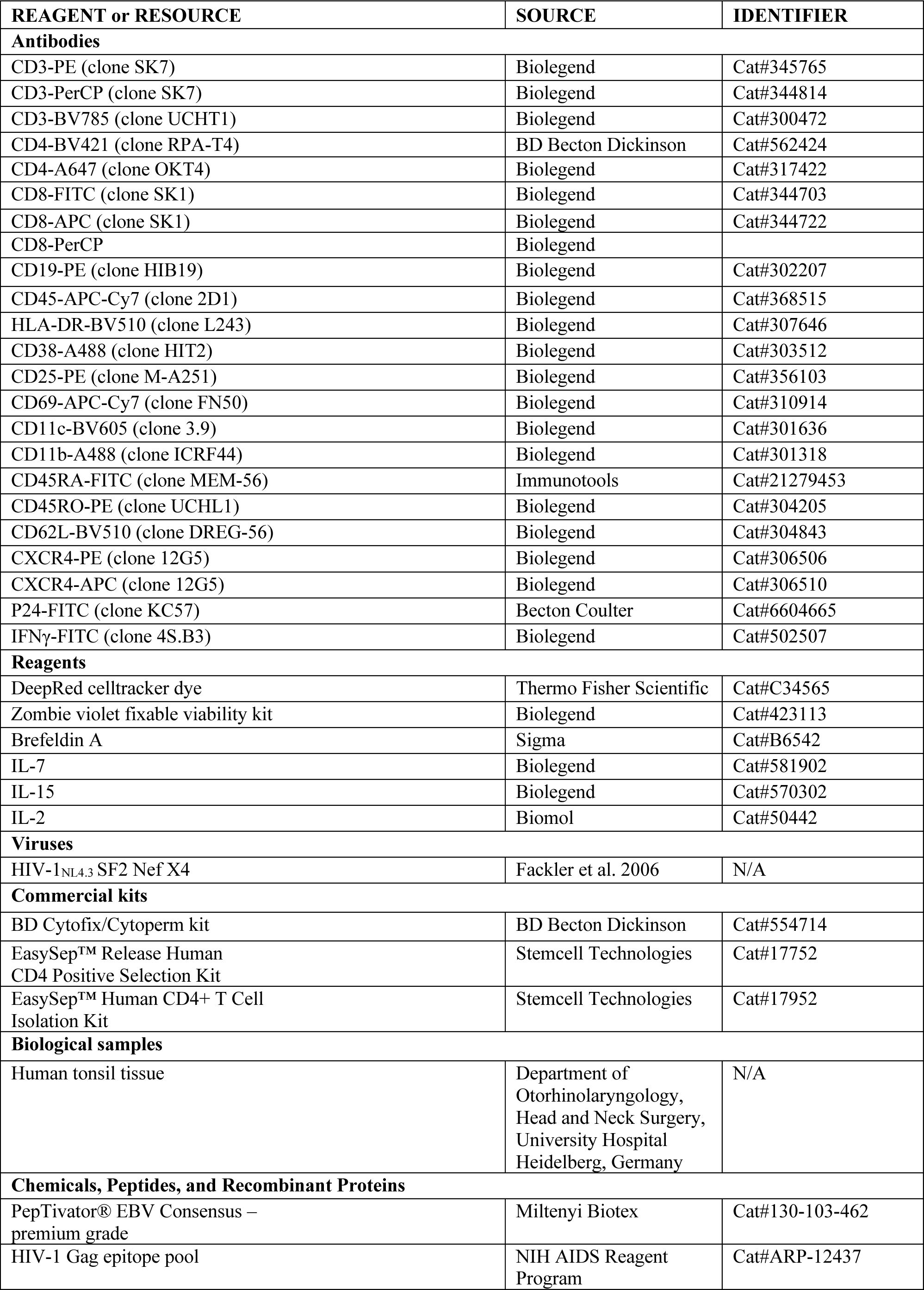
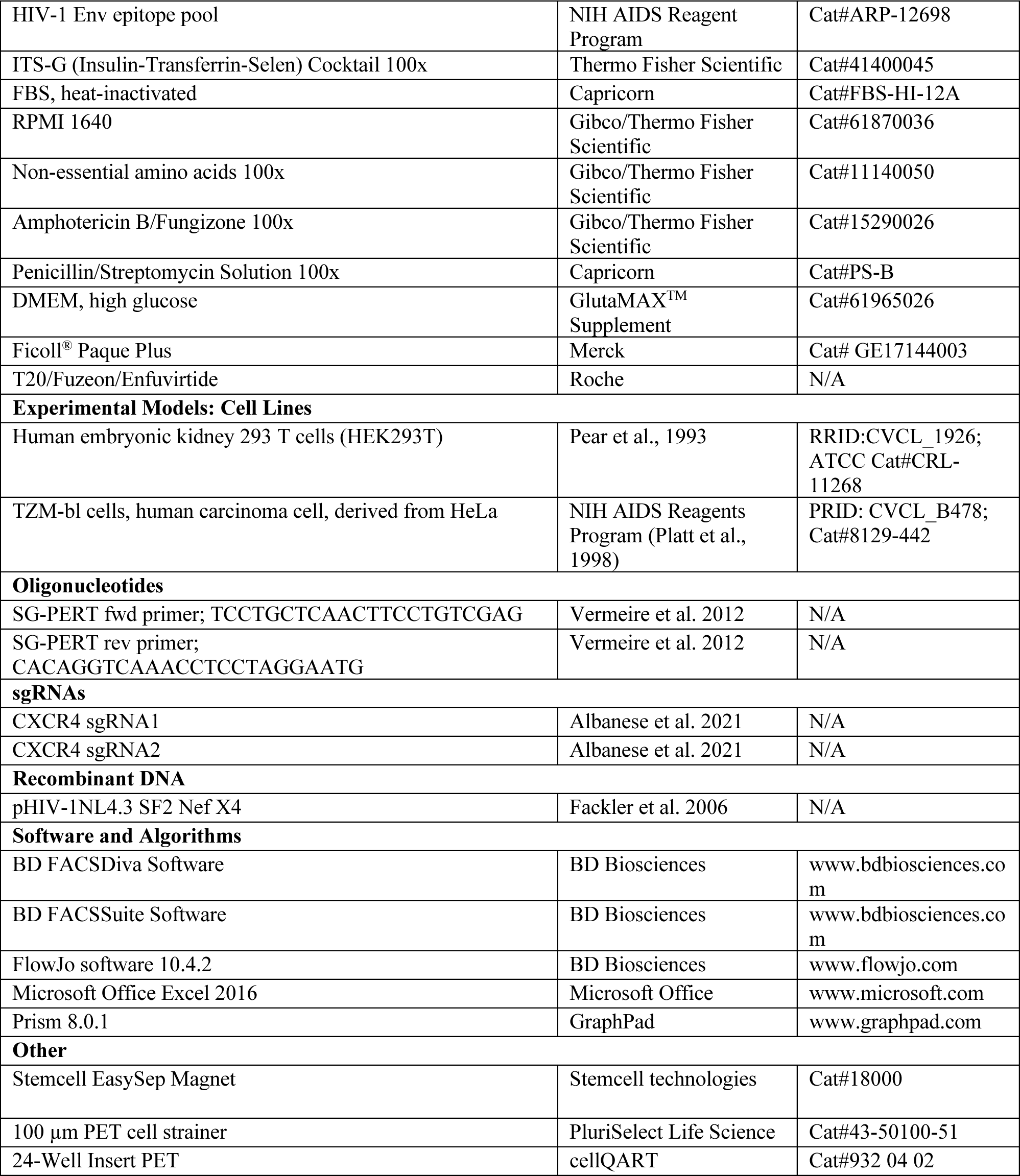

## Supplementary material

**Fig. S1.**
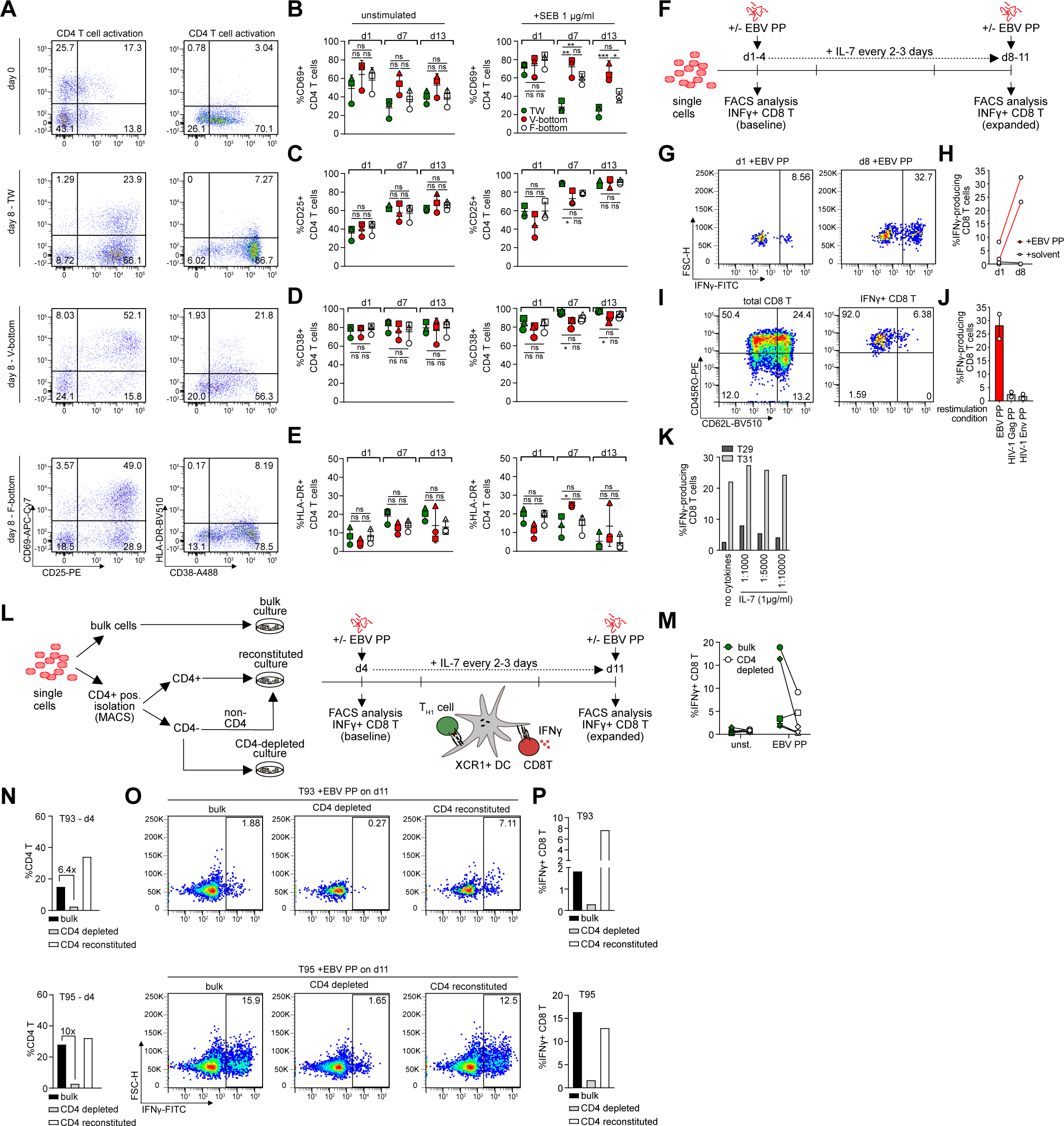
(Related to Fig. 1 and Fig. 2). (A) Representative dot plots of CD4 T cell surface activation marker expression showing frequency of cells positive for CD25 and CD69 (left panel) or CD38 and HLA-DR (right panel) on day 0 and day 8 of culture as indicated with (+SEB 1 µg/ml) exogenous activation in transwell, V- or F bottom plate culture. (B)-(E) Quantification of A without (left panel) or with (right panel) exogenous activation for three tonsils in TW (green), V-bottom (red) or F-bottom (white) setups. Shown are means with SD. Each symbol represents one donor. Statistical significance was assessed by one-way ANOVA (ns, p>0.05; *, p<0.05; **, p<0.01; ***, p<0.001; ****, p<0.0001). (F) Schematic of the experimental set-up for assessment of EBV-specific memory CD8 T cell reactivation in tonsil culture as in Fig. 2G. (G) Representative dot plots of tonsil CD8 T cells showing IFNγ-production upon EBV peptide stimulation on day 1 (baseline) and day 8 (expanded). (H) Quantification of G for two tonsils, showing increase of EBV-specific CD8 T cells that are identified by IFNγ-production over the culture period. Solvent control is water. (I) Representative dot plots of tonsil CD8 T cells after 7 days of culture with EBV peptide pool, showing typical distribution of memory subsets among total or IFNγ+ tonsil CD8 T cells measured by CD45RO and CD62L expression upon restimulation with EBV peptide pool, confirming an effector memory phenotype of the reactivated CD8 T cell response. (J) Quantification of IFNγ-production among CD8 T cells after 7 days of culture with EBV peptide pool and upon restimulation with EBV peptide pool or unspecific/background IFNγ-production upon restimulation with HIV-1 Gag and HIV-1 Env peptide pools, validating EBV-specificity of the expansion and reactivation process. (K) Quantification of IFNγ+ CD8 T cells upon restimulation with EBV peptide pool after 7 days of culture with EBV peptide pool and additional supplementation of IL-7 at indicated concentrations for two tonsils, respectively. (L) Schematic of the experimental set-up for assessment of CD4 T cell help to EBV-specific memory CD8 T cell reactivation. Briefly, tonsil single cell suspensions were seeded as bulk cultures or subjected to CD4 T cell positive isolation and the CD4-depleted fraction was seeded with or without reconstitution of CD4 T cells. The cultures were stimulated with EBV peptide pool on day 4 and expansion was assessed 7 days later. (M) Quantification of IFNγ+ CD8 T cells on day 11 after 7 days of culture with (EBV PP) or without (unst) EBV peptide pool for bulk cultures (grey) or CD4-depleted cultures (white). Each symbol represents one donor. (N) Frequency of CD4 T cells among tonsil cells upon initial peptide stimulation on day 4 for bulk, CD4-depleted and CD4-reconstituted cultures for two tonsils, respectively. (O) Dot plots showing IFNγ-production of tonsil CD8 T cells upon EBV peptide pool restimulation on day 11 after 7 days of culture with EBV peptide pool for bulk, CD4-depleted and CD4-reconstituted cultures for the two tonsils from N. (P) Quantification of O for the two tonsils.

**Fig. S2.**
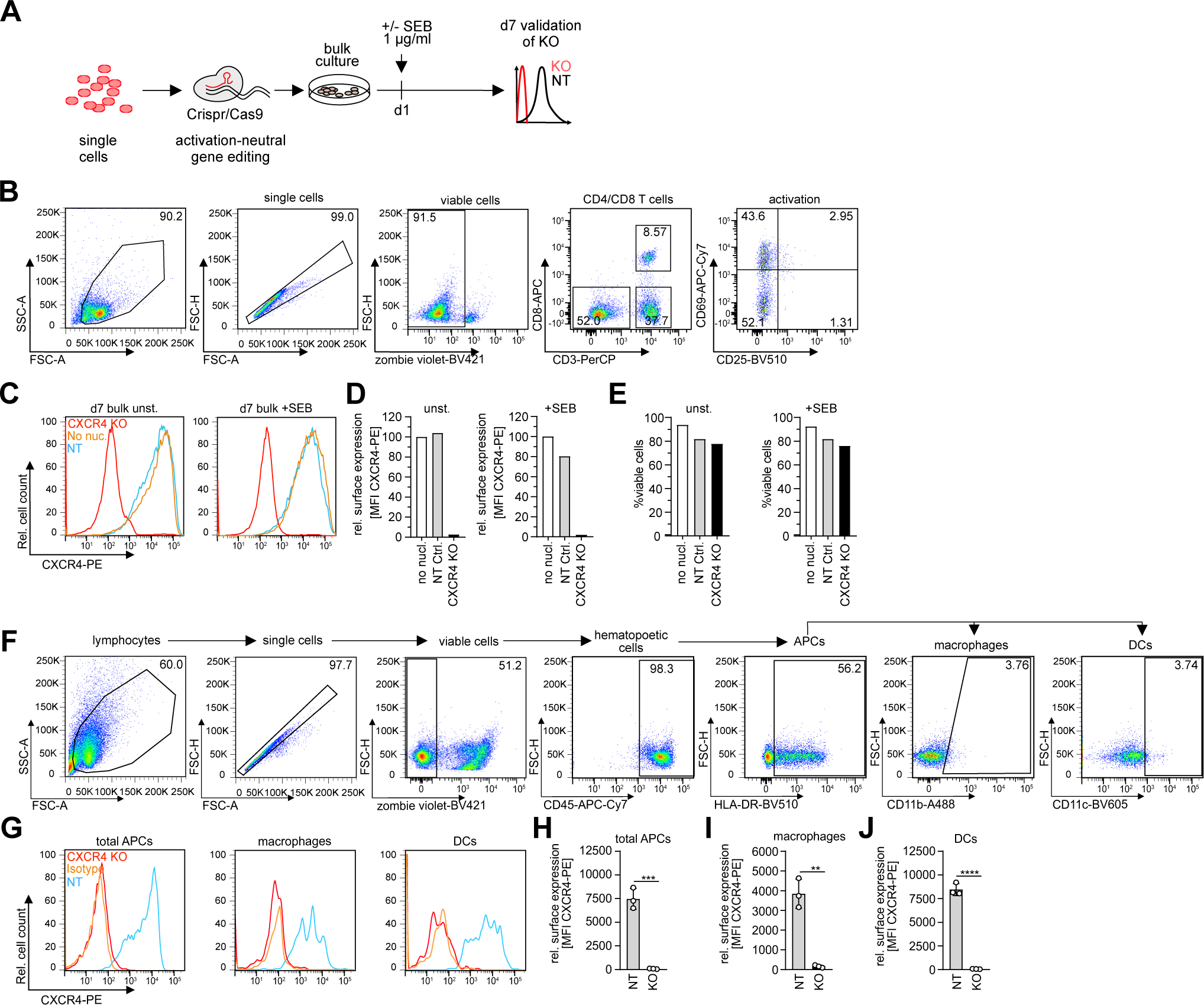
(Related to Fig. 3). (A) Schematic of the experimental set-up. (B) Representative dot plots showing gating strategy and typical activation phenotype of tonsil CD4 T cells upon seeding in the absence of exogenous activation, measured by CD25 and CD69 expression. (C) Histograms showing CXCR4 surface expression on CD4 T cells in tonsil bulk cultures without (left) or with (right) SEB stimulation at 1 µg/ml on day 7 post nucleofection. SEB was added 1 day post nucleofection. Superimposed are histograms for the non-targeting control (NT) and no nucleofection (no nucl.) condition. (D) Quantification of C for one tonsil without (unst.) or with (+SEB) exogenous activation on day 7 post nucleofection. (E) Frequency of viable CD4 T cells for conditions in C. (F) Representative dot plots and gating strategy for identification of myeloid subsets among tonsil cell cultures. (G) Representative histograms showing CXCR4 surface expression on total APCs, macrophages or dendritic cells (DCs) on day 7 post nucleofection with CXCR4-targeting RNP (CXCR4 KO) with superimposed histograms for the non-targeting RNP (NT) condition and isotype control. (H)-(J) Quantification of G for NT or CXCR4 KO in total APCs, macrophages or DCs. Each dot represents one tonsil. Shown are means with SD. Statistical significance was assessed by unpaired t-test (ns, p>0.05; *, p<0.05; **, p<0.01; ***, p<0.001; ****, p<0.0001).

**Fig. S3.**
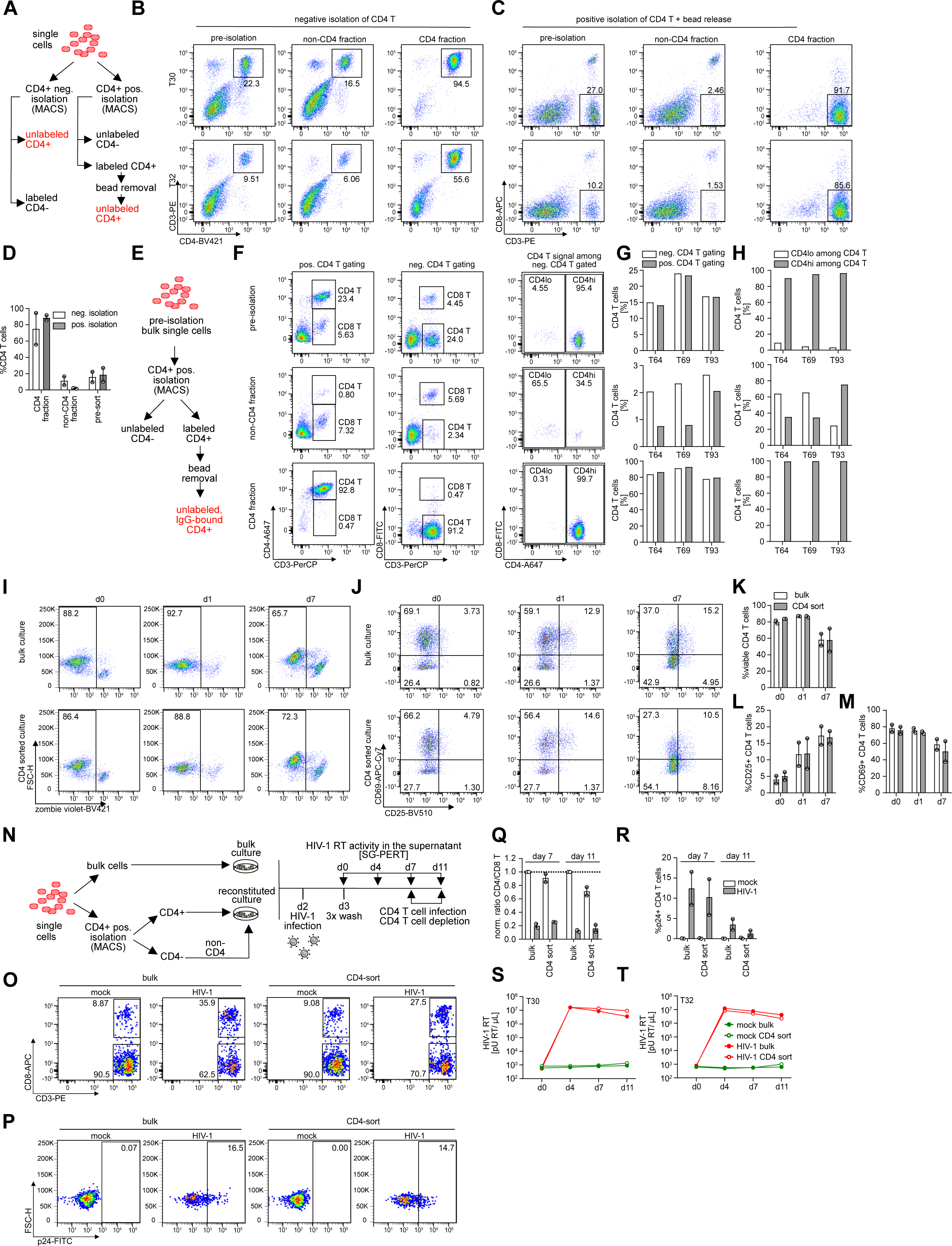
(Related to Fig. 4). (A) Schematic of the experimental set-up for CD4 T cell positive or negative isolation for comparison of sorting efficiency with indicated fractions expected from the two procedures. (B) and (C) Dot plots of CD4 negative (B) or positive (C) isolation fractions before (pre-isolation) and after sorting (non-CD4 fraction and CD4 fraction) for two tonsils. Indicated are frequencies of CD4 T cells based on CD3 and CD4 surface staining for negative isolation (B) or CD3 and CD8 surface staining for positive isolation (C). (D) Quantification of B and C for the two tonsils. Shown are means with range. Each dot represents one tonsil. (E)-(H) Assessment of the residual CD4 T cells left in the non-CD4 T cell fraction after positive isolation. (E) Schematic of the experimental set-up for positive isolation, indicating the resulting populations of antibody-bound, bead-released CD4 T cells and unlabeled non-CD4 T cells. (F) Representative dot plots showing tonsil cells pre-isolation, the non-CD4 T cell fraction post-isolation and CD4 T cell fraction post-isolation stained with CD3, CD8 and CD4 antibodies but showing either positive CD4 T cell gating based on CD3 and CD4 staining (left panel) or negative CD4 T cell gating based on CD3 and CD8 staining (middle panel). The right panel shows CD4 signal among negatively gated CD4 T cells based on CD3 and CD8 staining in the indicated fractions pre- and post-isolation. The CD4 antibody that was used in this experiment is the OKT4 clone which has a distinct binding site from the CD4-binding antibodies used in the positive isolation procedure. (G) Quantification of left and middle panel in F, indicating frequency of CD4 T cells based on the two gating strategies in the same samples. (H) Quantification of the right panel in F, indicating true frequency of CD4-positive cells among negatively gated CD4 T cells. (I)-(M) Validation of the activation-neutral CD4 T cell positive isolation procedure. (I) and (J) Representative dot plots showing viability (I) or activation (J) by surface expression of CD25 and CD69 of CD4 T cells on day 0, day 1 or day 7 in untouched bulk cultures or after CD4 T cell positive isolation and reconstitution of the culture. (K)-(M) Quantification of I and J for two tested tonsils at the respective timepoints. Shown are means with range. Each dot represents one tonsil. (N) Schematic of the experimental workflow to assess impact of the CD4 positive isolation on HIV-1 infection dynamics. Briefly, tonsil cells were thawed and infected with HIV-1 NL4.3SF2Nef at 1.5x10^5^ BCU per 2x10^6^ cells one day after CD4 positive isolation and reconstitution or bulk culture seeding. On the following day (d0), cells were washed 3x and infection was assessed by analysis of CD4 T cell infection and depletion by flow cytometry analysis upon final harvest. Further, production of HIV-1 virions in the supernatant was measured by SG-PERT throughout the infection course. (O) and (P) Representative dot plots showing CD4 and CD8 T cells subgated from CD3 T cells (O) as well as p24 signal in CD4 T cells (P) in bulk or CD4 T cell-sorted mock or HIV-1 infected conditions, respectively on day 7 post infection. (Q) and (R) Quantification of O and P for two tonsils with the ratio of CD4 vs. CD8 T cells in Q normalized to the mock condition for the respective bulk cultures. Shown are means with range. Each dot represents one tonsil. (S) and (T) HIV-1 RT activity in the supernatant of mock or HIV-1 infected bulk or CD4 T cell-sorted cultures at indicated timepoints for the two tonsils from O-R, respectively.

**Fig. S4.**
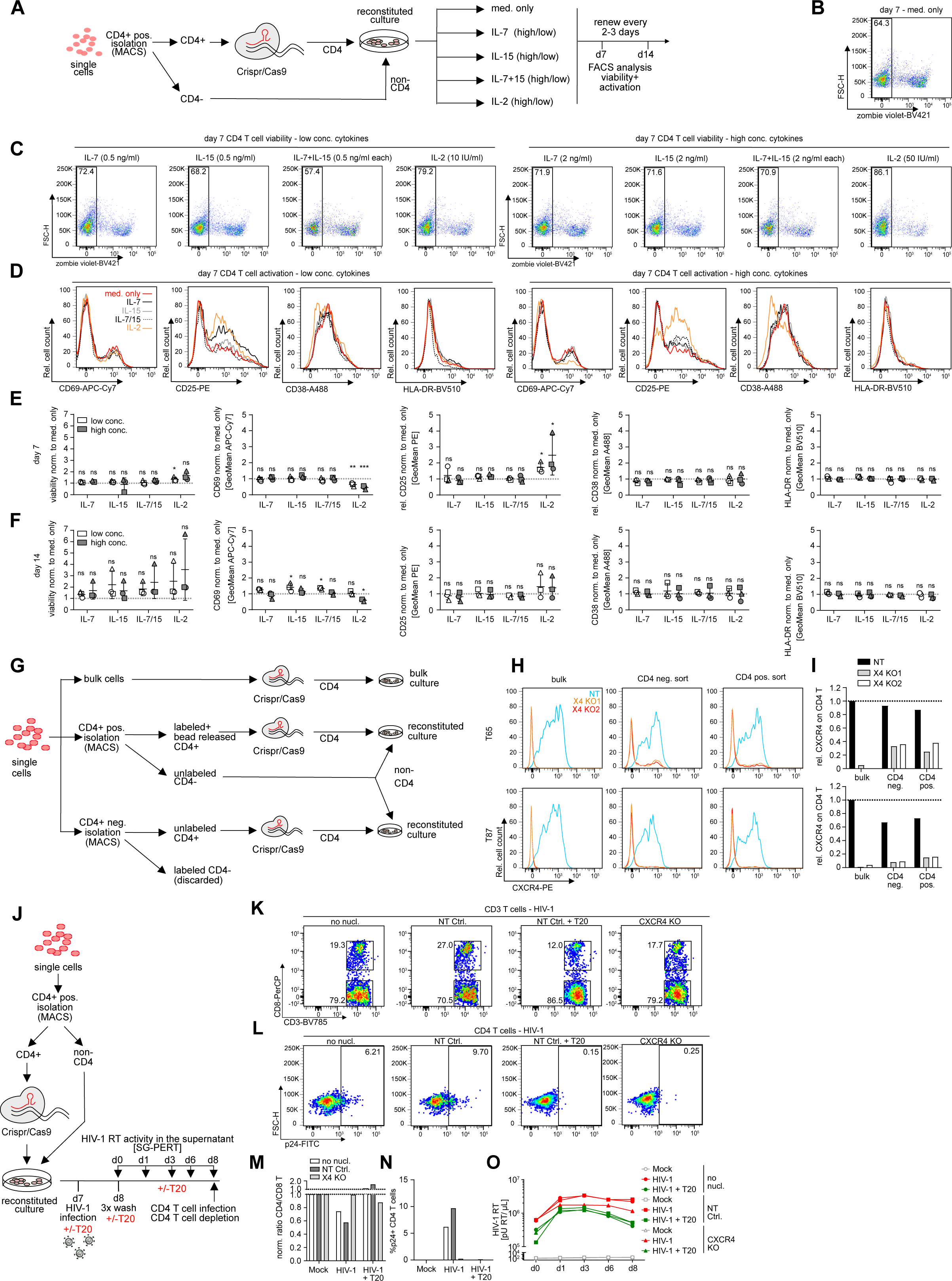
(related to Fig. 4). (A) Schematic of the experimental workflow. Briefly, tonsil cells were thawed, CD4 T cells were sorted, nucleofected with NT Ctrl. RNP and merged with non-CD4 T cells and cultured in transwell plates. Reconstituted cultures were kept in tonsil medium without further supplements (med. only) or with IL-7, IL-15 (separate or in combination) or with IL-2 with a low or high concentration as indicated. (B) and (C) Representative dot plots of the med. only (B) or cytokine-supplemented cultures with low (left panel) or high (right) concentrations on day 7 showing viability of the CD4 T cells based on zombie violet staining. (D) Representative histograms of CD4 T cells showing surface expression of activation markers CD25, CD69, CD38 or HLA-DR in low (left panel) or high (right panel) cytokine conditions. Superimposed are histograms for the med. only, IL-7, IL-15, IL-7+IL-15 or IL-2 cultures as indicated. (E) and (F) Quantification of D for day 7 (E) or day 14 (F) of culture. For each donor, values were normalized to the med. only condition. Each symbol represents one donor. Shown are means with SD. Statistical significance was assessed by one-way ANOVA (ns, p>0.05; *, p<0.05; **, p<0.01; ***, p<0.001; ****, p<0.0001). (G) Schematic of the experimental set-up for determining nucleofection efficiency in bulk or CD4 negatively vs. positively-isolated CD4 T cells. Briefly, tonsil cells were thawed and either nucleofected as bulk cultures or processed for CD4 negative or CD4 positive isolation with subsequent bead release. CD4 T cells resulting from the two separation procedures were nucleofected and merged with the unlabeled, CD4-depleted fraction resulting from the CD4 positive isolation while labeled non-CD4 T cells resulting from the CD4 T cell negative isolation were discarded. (H) Histograms showing CXCR4 surface expression on CD4 T cells in either bulk nucleofected cultures or CD4 negatively and CD4 positively-isolated and nucleofected populations on day 7 post nucleofection for two tonsils. Two individually prepared RNP complexes (X4 KO1 and X4 KO2) were tested side by side in each condition to test for variability of the complex quality in addition to CD4 T cell sorting strategy and to estimate overall robustness of the workflow. Superimposed are histograms for the respective non-targeting control (NT) condition. (I) Quantification of CXCR4 surface levels on CD4 T cells as shown in H for the two tested tonsils and the two RNP preparations normalized to the bulk NT values for each tonsil. (J)-(N) Validation of the inhibitory effect of CXCR4 KO in CD4 T cell sorted cells in the context of HIV-1 infection by comparison with T20 entry inhibitor of HIV-1. Briefly, CD4 T cells were sorted, nucleofected, merged with non-CD4 T cells and infected with HIV-1 after 7 days of culture in the presence or absence of T20 in transwell plates. (J) Schematic of the experimental set-up to compare efficiency of CXCR4 KO with chemical entry inhibition in the context of HIV-1 infection. Briefly, tonsil CD4 T cells were positively isolated, nucleofected and after 7 days, infected with HIV-1 NL4.3SF2Nef at 1.5x10^5^ BCU per 2x10^6^ cells in the presence or absence of T20/Enfuvirtide throughout the culture period. (K) and (L) Dot plots showing CD4 and CD8 T cells subgated from CD3 T cells (K) as well as p24 signal in CD4 T cells (L) on day 8 post infection in the non-nucleofected condition or NT Ctrl (untreated or with T20 treatment) or CXCR4 KO condition as indicated. (M) and (N) Quantification of K and L for one tonsil with the ratio of CD4 vs. CD8 T cells in M normalized to the mock condition for the respective conditions. (O) HIV-1 RT activity in the supernatant of mock or HIV-1 infected CD4 T cell-sorted cultures for the different conditions at indicated timepoints for one tonsil.

